# Morphological and biochemical evaluation of tolerance to water deficit identifies *SlCLE9* as a novel regulator of drought stress

**DOI:** 10.64898/2026.09.22.753559

**Authors:** Melek Ekinci, Selda Ors Cirik, Hong Le, Advait Anilal, Jordan Oxendine, Ruiting Wang, Shane Conner Grant, Rakesh K Upadhyay, Charles A. Seller, Daniel Rodriguez-Leal

**Affiliations:** Department of Horticulture, Faculty of Agriculture, Atatürk University, Erzurum 25200, Türkiye; Department of Agricultural Structures and Irrigation, Faculty of Agriculture, Atatürk University, Erzurum 25200, Türkiye; TomatoLab. Department of Plant Science and Landscape Architecture. University of Maryland, College Park, MD, 20742, USA; Department of Natural Sciences, Bowie State University, Bowie, MD, 20715. USA; Department of Cell Biology and Molecular Genetics. University of Maryland, College Park, MD, 20742, USA

**Keywords:** tomato, drought tolerance, *SlCLE9*, CRISPR/Cas9, heirloom tomato

## Abstract

We performed a screening to evaluate the responses to water deficit before the flowering stage in a panel of 31 genotypes composed of one commercial hybrid, one wild species, multiple ex-PVP (vintage) lines, several commercial heirlooms from the Northeast USA, Mexican landraces from hot and humid environments and genome edited lines with altered flowering and inflorescence traits. We characterized 18 different morphological, physiological and biochemical traits to assess the response to 40% water deficit. We observed diverse morphological, physiological and biochemical responses across the 31 genotypes screened, suggesting the presence of useful variation for breeding for drought tolerance in cultivated tomato. Interestingly, some heirlooms exhibited superior performance under drought, suggesting they can be used as donors to discover novel genes involved in drought responses. Unexpectedly, we found mutants in *SlCLE9*, a paralog of the stem cell regulator *SlCLV3* known to co-regulate meristem proliferation in tomato, exhibited enhanced drought tolerance. Notably, the *slcle9* mutant also conferred drought tolerance when used as rootstock with grafted scions from our reference cultivar M82, suggesting *SlCLE9* influence root architecture and drought response. Consistent with this, we found *SlCLE9* is transcriptionally upregulated in roots and leaves during drought stress, and exogenous application of SlCLE9 modifies root growth and response to drought. Altogether, our screening uncovered new potential trait donors for breeding against drought stress and describes a novel role for *SlCLE9* in drought tolerance.

**Main Conclusion:** We have identified several tomato genotypes as potential donors for drought tolerance and discovered a novel role for the gene *SlCLE9* in drought tolerance in tomato.

## Introduction

The main goal of plant breeding is to improve crops for food, feed and fuel. This process is achieved by a continuous cycle of selection for improved characteristics in breeding populations (Zamir 2001; He et al. 2024). However, breeders are often challenged by the lack of sources for genetic variation for multiple traits, the lengthy cycle of crops from seed to seed and the challenge of stacking multiple favorable loci for polygenic traits (Zamir 2001). In the past, enhancing genetic and phenotypic variation in crops has relied on intercrossing with wild relatives to introduce ‘‘exotic’’ allelic diversity, creating novel alleles by random mutagenesis, and genetic engineering (Zamir 2001). Remarkably, new technologies such as genome editing bring great promise for enhancing the genetic pool used for plant breeding, particularly for specialty varieties and for complex traits that may require simultaneous modification of several loci to confirm their association or to engineer novel variation (Nerkar et al. 2022).

Drought stress affects most agricultural areas in the world. Low rainfall rate, salinity, high and low temperatures, and high light intensity are among the factors contributing to drought stress. Drought stress causes significant changes in the morphological, physiological, biochemical and molecular properties of plants (Farooq et al. 2009; Bhargava and Sawant 2013). Specifically, drought stress affects membrane permeability, photosynthetic activity, oxidative metabolism, and ultimately leads to xylem embolism and cell death (Fahad et al. 2017). Therefore, plants have evolved physiological and developmental mechanisms to reduce water loss during drought stress, including stomatal closure and inhibition of photosynthesis, osmotic adjustment, decreased leaf area and stem elongation as well as modifications of the root system’s architecture (Chaudhry and Sidhu 2022). The ability of a plant to survive, adapt and recover from drought under wild conditions or in an agricultural setting is considered a complex trait, likely regulated by multiple genes influencing multiple metabolic processes (Conti et al. 2023). Multiplex genome editing represents a promising strategy to introduce drought tolerance in crops but requires precise identification of target genes that show potential to regulate drought tolerance (Lorenzo et al. 2023). Tomato (*Solanum lycopersicum* L.) is one of the world’s most-consumed vegetable crops (Anyaoha et al. 2023). It is a basic ingredient in a large variety of raw, cooked or processed foods and is produced globally for both local consumption and as an export crop. According to recent estimates from FAOSTAT27, the biggest producers of tomato worldwide are China, Turkey, India, USA and Italy (FAOSTAT 2023).

Tomatoes are susceptible to several abiotic stresses, including drought (Conti et al. 2023). Therefore, breeding for drought tolerant cultivars has become a priority in many global markets and cultivation types (open field, protected and urban agriculture) (Taheri et al. 2022). Although previous efforts have been made to develop an improved plant architecture by modifying genes controlling height, flowering or root growth, there has been relatively little efforts in breeding drought tolerance for controlled indoor or urban production in tomato, which is yet an underserved market particularly relevant for local agriculture in urban areas in the USA (Conti et al. 2023; Oxendine et al. 2026). This is particularly relevant for specialty varieties like heirloom tomatoes, which are known for their superior taste but lack of improved tolerance to abiotic stresses (Mid-Atlantic Commercial Vegetable Production Recommendations | University of Maryland Extension). In contrast, much has been done around improving cultivation conditions (Aggarwal et al. 2024). Therefore, there is a critical need to develop improved cultivars suitable for urban agriculture that retain superior quality and tolerance to abiotic stresses such as drought (Aggarwal et al. 2024). Tomato is a global crop subject to different growing conditions, and several seed banks and collections are available for screening for tolerance to abiotic stresses.

In this study, we took advantage of these resources to evaluate the range of phenotypic variability to water deficit in tomato. Screening a collection of tomato varieties, including one wild species, several landraces from hot and dry environments in Mexico, vintage varieties including heirloom tomatoes grown in the Northeast USA, and genome-edited lines with mutations in plant architecture and flowering time genes, we found diverse responses to water deficit. We observed major differences in biomass allocation or their biochemical and physiological responses. We identified several lines with enhanced drought tolerance and thus with potential to be used as donors in breeding programs. Particularly, the heirloom variety Prudens purple and the Mexican landrace Guaco, which comes from a very hot and humid area in the State of Sinaloa, Northwest Mexico showed the best overall tolerance to drought. Notably, we found the mutant *slcle9*, which encodes for the closest homolog of the tomato shoot meristem regulator *CLAVATA3* exhibited optimized root biomass allocation, carbon assimilation and water use efficiency under drought, suggesting *SlCLE9* has a novel role in plant development and stress resilience in the vegetable crop tomato.

## Materials and Methods

### Growth conditions and selection of cultivars for drought screening

We worked with selected genotypes from a small collection of 60 lines comprising one wild species, several landraces, ex-PVP (vintage) and heirloom varieties, and genome-edited mutants showing differences in plant architecture and flowering time. These materials were obtained from the Tomato Genetics Resource Center (TGRC), from commercial vendors (Johnny’s Seeds, Annie’s Heirlooms) or donated by Prof. Zachary Lippman (CSHL). We focused our screening on a subset of vintage, heirlooms and genome edited lines that we considered more relevant for the urban agriculture sector in Maryland based on whether they are commercially relevant heirlooms or exhibit interesting traits (early flowering, semi-determinate growth, potential resistance biotic and abiotic stress). We narrowed down the list to 31 genotypes from this collection (**Supplemental Table 1**). Tomato seeds were germinated in 96-inset flats using a germination soil mixture (SunGro). Individual seedlings were transplanted into 6-inch pots 3 weeks after germination in potting soil (professional growing mix; Canadian sphagnum peat moss, perlite, dolomite lime, long-lasting wetting agent, resilience, Sunshine® and 10 g of Osmocote slow-release fertilizer). Greenhouse experiments were performed during Spring 2025, under long days (16 hr), with day temperatures of 25°C and night temperatures of 18-20°C.

### Water deficit and drought experiments

For water deficit treatments, greenhouse grown plants were irrigated at 100% of the field capacity (full irrigation-control) and 60% or 50% of the field capacity (40% or 50% water deficit, respectively). The amount of water given to pots was calculated according to the field capacity of the control pots. For this purpose, the amount of water added to control and treatment pots was determined according to volume-based moisture values measured with a moisture meter (TDR, Spectrum Technologies, Inc.) every 2-3 days in the morning. The study was set up in a random plot trial design with 3 replications and 2-6 plants in each replication.

Drought experiments with complete water withholding comparing M82 and *slcle9* plants and grafted individuals were performed under greenhouse or growth chamber conditions. Briefly, M82 and *slcle9* seeds were germinated in 96-inset flats using germination soil mix. Individual seedlings were transplanted into 6-inch pots with potting soil 3 weeks after germination. For grafting plants, selected plants were cut using a clean, sharp knife, approximately 2–3 cm above the cotyledons at a 45° angle to increase the contact surface. We used a grafting clip to secure the scion and rootstock at the graft junction, followed by inserting a straw or similar support next to the graft to help stabilize the plant. Grafted plants were maintained at high humidity and in the dark for 5-7 days. After that, plants were placed in a humid environment and exposed to a 16-hour light/8-hour dark photoperiod with a light intensity of approximately 200 µmol m^2^ s^-1^. After two weeks, grafted plants were used for drought experiments under greenhouse conditions. For each control (full irrigation) and treatment (no irrigation for ∼10 days), plants were placed in trays with 8 plants each. After the 11^th^ day of the experiments, data was collected on different morphological traits. Control plants were watered with 50 mL of deionized water every day, while drought treatments consisted of withholding water for 8-11 days until wilting symptoms were observed. This experiment was performed using 8 plants per genotype per condition (M82 vs *slcle9*, control vs treatment, N = 32). We collected gas exchange (*gsw*), carbon assimilation (*A*) and intrinsic Water Use Efficiency (*iWUE =* A/gsw) data from the expanded 4^th^ leaf of three independent plants per genotype over a time course of 35 minutes (data capture every 5 seconds) per plant using a Licor-6800 device (LI-COR).

To evaluate genotype performance, stability, and stress susceptibility across the screening population, two complementary stress indices known as the Stress Susceptibility Index (SSI) and the Stress Tolerance Index (STI) were calculated for all evaluated traits using paired mean values under well-watered control (*Y_p_*) and water deficit (*Y_s_*) conditions. The Stress Susceptibility Index (SSI) (Fischer and Maurer 1978) was calculated according to quantify relative performance reduction under drought compared to non-stress conditions:

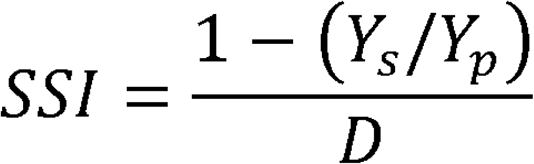

Where:

*Y_s_* is the mean performance of a given genotype for a specific trait under water deficit (40% water deficit).

*Y_p_* is the mean performance of the same genotype for the trait under well-watered control conditions (100% field capacity).

*D* represents the overall stress intensity index across the entire population, calculated as:

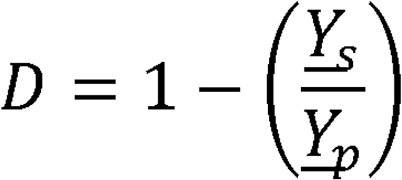

Where *Y_s_* and *Y_p_* are the overall population means across all genotypes under stress and non-stress conditions, respectively. Lower SSI values (<1) reflect lower trait reduction and higher stress stability under water deficit. The Stress Tolerance Index (STI) (Fernandez 1992) was calculated to identify genotypes that achieve high performance under both non-stress and stress conditions:

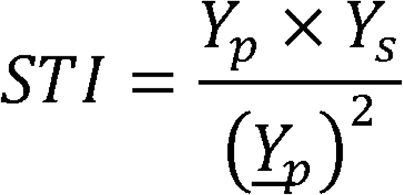

Where Y_p_ and Y_s_ represent the genotype-specific trait means under control and stress conditions, respectively, and *Y_p_* is the overall population mean under well-watered control conditions. Higher STI values indicate superior performance potential and trait maintenance across both environments. To assess multi-trait performance, traits were categorized into three functional groups: vegetative biomass (PH, SD, LA, PFW, RFW, PDW, RDW), physiological parameters (LRWC, EC, Chl a, Chl b, Total Chl, Car), and biochemical responses (H_2_O_2_, MDA, SOD, CAT, POD). Category-specific STI and SSI scores were computed as the arithmetic mean of the individual trait indices within each respective functional group. For multi-trait heatmap visualizations and biplot comparisons, raw index values were column-standardized to *Z*-scores (µ = 0, σ = 1) to facilitate direct cross-trait comparisons.

### Determining the effect of water deficit in tomato development

For the 40% and 50% water deficit experiment, we measured plant height (PH), stem diameter (SD), plant fresh weight (PFW), plant dry weight (PDW), root fresh weight (RFW) and root dry weight (RDW) of all plants in all replicates of the control and treatments at the end of each experiment 40 days after transplanting. The plant and root material was kept at 67 °C for 48 h for dry weight measurements. Plant total leaf area (LA) was determined by the leaf area meter (LI-3100 Area Meter; Li-Cor, USA). Electrolyte leakage (EL) and leaf relative water content (LRWC) of the plants were calculated as previously (Yildirim et al. 2021). The plant samples were kept at -86 °C until analysis. Before analysis, tissue samples were ground in liquid nitrogen using a mortar with a pestle and dissolved in acetone. The concentrations of chlorophyll a (Chla), chlorophyll b (Chlb), carotenoid and total chlorophyll (Total Chl; calculated with a + b) were measured spectrophotometrically at 645, 663 and 450 nm wavelengths determined and calculated as milligrams per gram.

For lipid peroxidation (MDA) analysis the absorbance of the supernatant obtained was measured at 400, 500, and 600 nm, and the MDA concentration was determined from the absorbance using an extinction coefficient of 155 mmol l^−1^cm^−1^ (Shams et al. 2019). Hydrogen peroxide (H_2_O_2_) was determined as previously (Velikova et al. 2000). The absorbance of the supernatant was measured at 390 nm and the content of H_2_O_2_ calculated by comparing it with a standard calibration curve made using different concentrations of H_2_O_2_. To measure antioxidant enzyme activities, we homogenized fresh leaf samples in the extraction solution following previously published protocols (Angelini and Federico 1989; Angelini et al. 1990). The obtained supernatant was used to determine enzyme activities of Superoxide dismutase (SOD), catalase (CAT) and peroxidase (POD) using a spectrophotometer with wavelengths of 560 nm, 240 nm and 436 nm, respectively (Angelini et al. 1990; Abedi and Pakniyat 2010).

### *In vitro* analysis of drought/osmotic stress

For *in vitro* osmotic/drought stress experiments, we adapted a published protocol for Arabidopsis seeds (Verslues et al. 2006). Briefly, we sterilized seeds of M82 and *slcle9* by first washing them in sterile water and 0.1% Tween 20 with vigorous shaking for 3 min, subsequently washing them in 70% ethanol and then in 2% bleach under constant shaking for 10 min. Sterilized seeds were germinated in MS plates (1.2 g/L MES, 2.2 g/L MS salts, and 15 g/L agar, adjusted to pH 5.7). Once the radicle emerged (∼5 mm length), seedlings were moved to control plates with the same media (no PEG) or plates containing polyethylene glycol (PEG-8000, Fisher). The cooled PEG (250 g/L) solution was poured onto the solidified agar plates at a PEG-to-agar ratio of 3:2 to achieve a water potential of -0.5MPa. The plates were left overnight (approximately 12–15 hours) at room temperature to allow the PEG solution to diffuse into the agar, after which the excess PEG solution was discarded. Six to eight seeds were placed in a straight horizontal row on each plate, and the plates were positioned vertically to ensure the roots grew straight downwards. After 6-8 days, data was collected on root length and number of lateral roots. Three technical replicates (3 plates with 6-8 seedlings) were included for each genotype and condition. For *in vitro* peptide treatments, we added a synthesized SlCLE9 peptide (SlCLE9p: REAPMSPDPLHH; Genscript) to both MS and MS+PEG plates to reach a concentration of 10 µM SlCLE9p. Negative controls consisted of plates without the peptide, and with or without PEG. Three technical replicates (3 plates with 6-8 seedlings) were included for each genotype and condition.

### RNA extraction and expression analysis

RNA was extracted from leaves and roots of plant samples before (Day 0) and after (Day 11) water withholding from the growth chamber drought experiment. Briefly, 150 mg of tissue was collected, and tissue samples were frozen in liquid nitrogen and ground using a GenoGrinder HG-600 (Cole Palmer). RNA was extracted using the Invitrogen PureLink RNA mini kit (ThermoFisher) and treated with DNase I (ThermoFisher) following manufacturer’s instructions. First-strand complementary DNA synthesis (cDNA) was made from 2 µg of RNA using the iScript Advanced cDNA synthesis Kit and SsoAdvanced universal SYBR Green SuperMix reagents. The CFX-Opus real-time PCR detection system was used for gene expression quantitation (Bio-Rad, Hercules, California, USA). Gene expression was derived according to the 2-ΔΔCT method (Livak and Schmittgen 2001). Quantitative reverse-transcription (qPCR) was performed on three biological and three technical replicates per genotype, condition and tissue using gene-specific primers against *SlCLE9* and 17 different marker genes associated with drought responses in tomato (Upadhyay et al. 2021). The gene *ADP RIBOSYLATION FACTOR* (*Solyc01g008000*) (Duan et al. 2022) was used as reference for all qPCR experiments. See **Supplemental Table 9** for the list of all primers used in this study.

### Statistical analysis

Statistical analysis on drought experiments under greenhouse conditions was performed by ANOVA using SPSS and the difference between the means was determined using the Duncan multiple comparison test. Linear Mixed Models (LMM) and a Negative Binomial Generalized Linear Model (GLM) were implemented for Licor-6800 data and data collected from *in vitro* experiments using RStudio scripts. qPCR data were analyzed following the methodology previously published protocols (Livak and Schmittgen 2001; Yuan et al. 2006) using scripts in RStudio. Relative gene expression was calculated using the delta-delta Ct method with the gene *ADP-Ribosylation Factor* (*Solyc01g008000*) as the internal reference. Factorial ANOVA was implemented using a linear model (*lm*), and *post-hoc* pairwise comparisons between genotypes and treatment conditions within each tissue type were evaluated using estimated marginal means within the package *emmeans* v1.10 (Lenth and Piaskowski 2026) in RStudio. *P-values* were adjusted for multiple testing using the False Discovery Rate (FDR) procedure.

## Results

### Plant growth and physiology are negatively affected by water deficit treatment in tomato

We evaluated a population of 31 genotypes for their response to water deficit during vegetative growth, prior to the transition to flowering as measured by the number of leaves (3-5) at the time of the experiment and the lack of a visible inflorescence. This collection comprised one wild species (*Solanum pimpinellifolium*), one commercial hybrid listed as drought tolerant (Mountain Lion F1), six ex-PVP (vintage) cultivars, six Mexican landraces from the States of Veracruz, Sinaloa and Mérida, eleven commercial heirlooms grown in the Mid-Atlantic USA and five different CRISPR/Cas9-generated mutants affected in flowering time and meristem development; which are traits that may influence or be influenced by drought (**Supplemental Table 1**) (Xu et al. 2015; Rodríguez-Leal et al. 2017; Soyk et al. 2017; Rodriguez-Leal et al. 2019). We evaluated a total of 18 morphological, physiological and biochemical traits under both full irrigation and 40% water deficit (**Figure 1**, **Supplemental Tables 2-7**). When all 31 genotypes were plotted using a PCA biplot, we observed a clear distinction between full irrigation and water deficit conditions, primarily driven by biomass traits (control) and stress related markers (water deficit treatment; **Figure 1a**). We observed significant differences in drought response among all these lines for most measured traits between control and treatments (**Figure 1b-i**, **Supplemental Tables 2-7** and **Supplemental Datafile 1**). We performed a Pearson’s correlation analysis to evaluate the relationships among the 18 evaluated traits across 31 genotypes under full irrigation and water deficit conditions. In both full irrigation and water deficit conditions, plant height (PH), stem diameter (SD), leaf area (LA), plant (PFW) and root fresh weights (RFW), and plant (PDW) and root dry weights (RDW) exhibited significant positive correlations (*r* > 0.30, *p* < 0.05). Specifically, leaf area (LA) showed strong positive associations with stem diameter (*r* = 0.67, *p* < 0.05 under full irrigation; *r* = 0.71, *p* < 0.05 under water deficit) and shoot fresh weight (PFW; *r* = 0.84, *p* < 0.05 and *r* = 0.82, *p* < 0.05, respectively). Chlorophyll a (Chl a), Chlorophyll b (Chl b), Total Chlorophyll (Total Chl), and Carotenoids (Car) were positively correlated with one another. As expected, the strongest relationship was observed between Chl b and Total Chl (*r* = 0.93, *p* < 0.05 under full irrigation; *r* = 0.64, *p* < 0.05 under water deficit), as well as Chl a and Total Chl (*r* = 0.84, *p* < 0.05 and *r* = 0.92, *p* < 0.05, respectively). Under water deficit, biomass parameters showed negative associations with photosynthetic pigments and carotenoids. Shoot dry weight (PDW) displayed moderate negative correlations with carotenoids (*r* = -0.46) and Chlorophyll a (*r* = -0.51, *p* < 0.05). Additionally, peroxidases (POD) showed significant negative correlations with morphological traits such as leaf area (*r* = -0.61, *p* < 0.05) and stem diameter (*r* = -0.44, *p* < 0.05) under drought. Antioxidant enzyme activity displayed condition-dependent relationships; for instance, superoxide dismutase (SOD) exhibited negative correlations with plant dry weights (PDW & RDW) under full irrigation (*r* = -0.10 and -0.48 *p* < 0.05, respectively) and retained a strong association with PDW under water deficit (*r* = -0.37, *p* < 0.05). Overall, this suggests that managing oxidative damage is a key component of drought tolerance.

**Figure 1.**
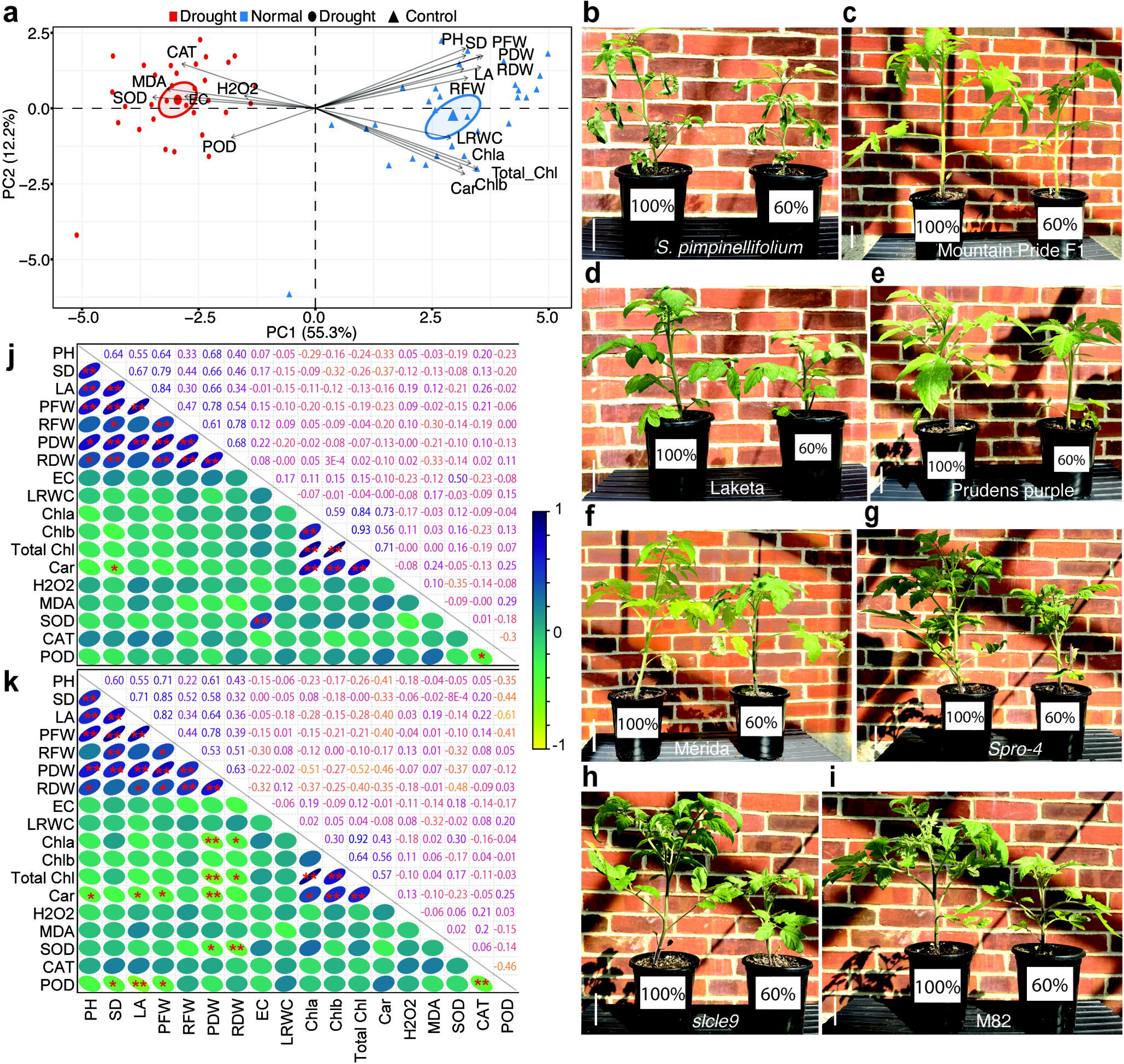
Multivariate phenotypic analysis and morphological responses of tomato genotypes under control and drought stress conditions. a, Principal Component Analysis (PCA) biplot showing the clustering of tomato genotypes and loading vectors for 18 traits across control (black triangles) and drought stress (black circles; 40% water deficit) treatments. Individual genotype responses under drought and normal conditions are color-coded in red and blue, respectively. PC1 and PC2 capture 55.3% and 12.2% of total phenotypic variance. b-i, Representative phenotypic responses of selected tomato genotypes grown under well-watered (100% field capacity; left pot) and drought stress (40% water deficit, right pot) conditions. b, *S. pimpinellifolium*. c, Mountain hybrid F1. d, Laketa. e, Prudens purple. f, Mérida. g, *spro-4*. h, *slcle9*. i, M82 indeterminate. j-k, Pearson correlation matrices showing pair-wise relationships among 18 morphological, physiological, and biochemical traits under well-watered control (100% field capacity; j) and drought stress (40% water deficit; k) conditions. Correlation coefficients (*r*) are displayed in the upper triangular matrix, while circle size, direction, and color intensity in the lower matrix represent the magnitude and direction of the correlation (scale bar from -1 to 1). Statistical significance is based on Benjamini–Hochberg false discovery rate (FDR) and annotated as * = *p* < 0.05 and ** = *p* < 0.01. Scale bars in b-i, 5 cm.

The differential response to water deficit was more pronounced in biomass accumulation. The greatest decrease in plant height (41%) and stem diameter (33%) was seen for the landrace Papantla de Olarte (LA4769). On the other hand, the heirloom Eva purple ball only exhibited a 5% decrease in plant height, and the heirloom Sunray tomato had a mild decrease in stem diameter (6%). Moreover, the ex-PVP Hotset (LA3320) exhibited the greatest decrease in leaf area (57%), while the wild species *S. pimpinellifolium* (LA1589) only exhibited 11% reduction under water deficit (**Supplemental Tables 2** and **8**). This was supported by an overall reduction of fresh and dry weights of shoots and roots (**Supplemental Tables 3** and **8**). Compared to full irrigation, the greatest decrease in PFW under water deficit was observed in the Hotset (57%), while the least decrease was found in *S. pimpinellifolium* (7%), followed by the hybrid commercial line Mountain Pride F1 (10%) and the heirloom Amana Orange (21%). For PDW, the greatest decrease was seen in the heirloom cultivar Amish gold (64%), whereas the CRISPR/Cas9 mutant *spro-4* (Rodriguez-Leal et al., 2017) exhibited a 26% decrease, followed by the heirloom varieties Carbon and Jubilee. For RFW, the Mexican landrace Papantla de Olarte exhibited the greatest decrease in RFW (65%), followed by the heirloom Eva Purple ball. In contrast, the ex-PVP Hotset showed a more robust response to water deficit by only reducing RFW by 12%, followed by the CRISPR/Cas9 mutant *slcle9*. For RDW, Hotset maintained higher biomass (7% decrease), followed by the Mexican landrace Papantla de Olarte (10% decrease) (**Figure 1a, Supplemental Tables 2, 3** and **8**). These results indicate the negative effects of water deficit for the morphological traits analyzed were not uniform across cultivars and that our germplasm collection exhibits wide phenotypic diversity to water deficit in cultivated tomato.

To complement our morphological assessment of the effects of water deficit, we also quantified several physiological parameters in our collection of 31 genotypes. Leaf Relative Water Content (LRWC) values decreased under water deficit, with the heirloom Sunray showing the largest decrease (-23%) and the heirloom Brandywine pink showing the smallest decrease among genotypes (-2%), followed closely by the ex-PVP Rehovot 13 (LA3129; -3%) and the Mexican landraces Mérida (LA1462) and La Biznaga (LA4752), the latter coming from arid or hot and humid environments (**Supplemental Tables 4** and **8**). Electrolyte leakage (EL) increased significantly under water deficit. Notably, the CRISPR/Cas9 mutant *spro-5*, which is known to increase inflorescence branching (Rodríguez-Leal et al. 2017) exhibited the largest increase of EC under water deficit (+285%) followed by the heirlooms Brandywine pink and Cherokee purple. On the other hand, the ex-PVP Laketa (LA0505) showed the smallest increase (+1%), followed by the heirloom Jubilee (+18%) and the commercial hybrid Mountain Pride F1 (+20%) (**Supplemental Tables 4** and **8**).

Total Chlorophyll (Total Chl) and Chl and Chl b values were also significantly affected under water deficit conditions. While the heirlooms Cherokee purple and Prudens purple showed the lowest decrease in total Chlorophyll content (-13 and -16%, respectively), the Mexican landrace Comala (LA2084) and the ex-PVP Globonnie (LA2802) were the most severely affected by water deficit (-74 and -70%, respectively). For carotenoid (Car) content, we observed that the Mexican landrace Xicotepec (LA4739) exhibited the lowest content under full irrigation conditions, however, it largely maintained this level under water deficit (10.72 mg/g vs 10.39 mg/g) showing the smallest decrease ratio among all lines (-3%). On the other hand, Globonnie again showed a high impact of water deficit (-56%) (**Supplemental Tables 5** and **8**). These results highlight contrasting drought-adaptation mechanisms, with some genotypes exhibiting higher membrane stability and photosynthetic capacity under moderate drought conditions.

To determine the biochemical response to water deficit, we also measured the levels of hydrogen peroxide (H_2_O_2_), Malondialdehyde (MDA) and the enzymatic activities of Catalase (CAT), Peroxidase (POD) and Superoxide Dismutase (SOD) (**Supplemental Tables 6-8**). We observed that H_2_O_2_ accumulation varied widely across our collection, with the heirloom Jubilee showing a +700% increase, followed by Comala (+650%) and Prudens Purple (+460%), while Brandywine pink exhibited the smallest increase (+8%), followed by the ex-PVP Break O’Day (LA1499). Interestingly, the highest levels of Peroxidase (POD) were observed in the CRISPR/Cas9 mutant *slclv3pro* and *sp5g* (+1525% and +1500%, respectively), while *slcle9* showed the lowest increase in POD activity (+40%), roughly half of its parental genotype M82 (+75%). These results revealed that some lines in our collection maintain biomass or trigger a strong physiological or biochemical response under water deficit conditions, suggesting the potential for several lines in our collection to serve as donors for improving different physiological responses to drought in cultivated tomato.

### Population clustering using drought tolerance indexes

To better understand the differences between the genotypes in our screening population, we ranked the genotypes based on their response to water deficit by calculating the Stress Susceptibility Index (SSI) (Azrai et al. 2025) and using calculated Z-scores (standard scores) to aggregate the SSI across multiples biomass parameters (**Figure 2a**). Our population displayed considerable diversity in trait-specific susceptibility across morphological, physiological, and biochemical parameters. The top-ranked, most tolerant genotypes *S. pimpinellifolium*, Mountain Pride F1, and Laketa consistently maintained low SSI Z-scores across biomass parameters (PH, SD, RDW, PDW, RFW, PFW, LA). In contrast, genotypes at the bottom of the ranking, such as *spro-5*, *slclv3pro*, Flora-Dade, and Break O’Day, demonstrated severe susceptibility. Interestingly, all genome-edited mutants share the same parental background (M82), but they did not necessarily rank closely to M82, suggesting these mutants have an altered response to drought. These results suggest functional trade-offs between biomass maintenance and physiological/biochemical adaptation across the population.

**Figure 2.**
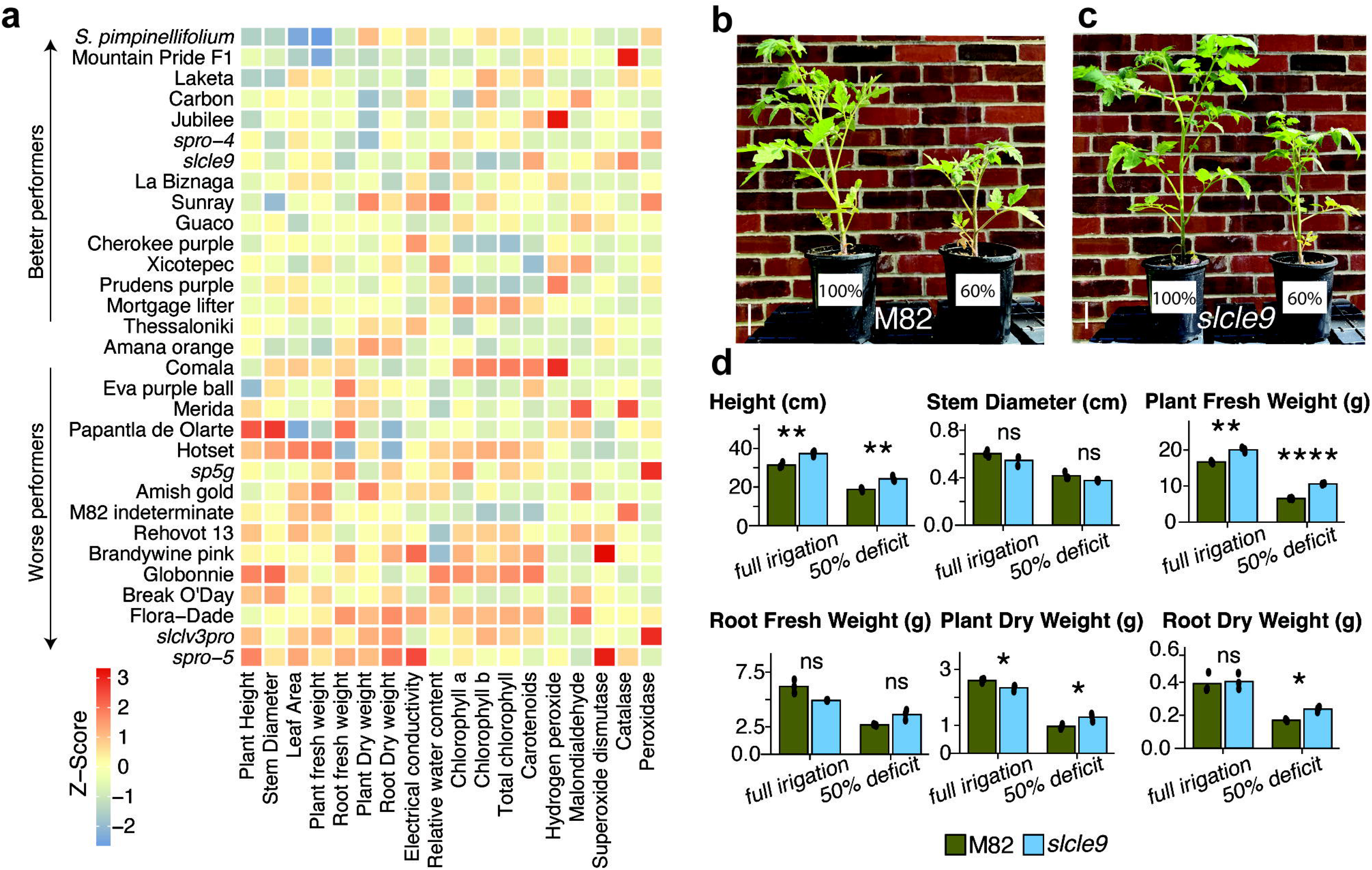
Ranking based on Stress susceptibility index and phenotypic comparison of M82 and the *slcle9* mutant under water deficit conditions. a, Heatmap displaying the column-standardized (*Z*-score) Stress Susceptibility Index (SSI) for 18 phenotypic parameters across 31 tomato genotypes, ranked top-to-bottom by biomass tolerance (lowest average biomass SSI at top). Color gradient denotes stress susceptibility relative to the population mean, where negative *Z*-scores (blue) represent lower susceptibility (higher stability) and positive *Z*-scores (red) indicate higher susceptibility. **b-c,** Representative pictures of M82 (b) and *slcle9* (c) plants grown under full irrigation (100% field capacity; left pots) and severe water deficit (50% field capacity; right pots). **d,** Quantitative measurements of biomass traits from M82 and *slcle9* plants under full irrigation and 50% water deficit conditions. Data represent means ± SD. Statistical significance between genotypes within each condition was determined by multifactorial ANOVA (ns = not significant, * *p* < 0.05, ** *p* < 0.01, **** *p* < 0.0001).

Highly biomass-tolerant accessions generally exhibited lower susceptibility Z-scores for oxidative stress indicators (H_2_O_2_, MDA) and membrane leakage (EC), suggesting better cellular protection during drought. Conversely, moderate to susceptible genotypes displayed variable enzymatic responses. For example, several genotypes positioned in the lower half of the ranking showed heightened SSI values for antioxidant enzymes (SOD, CAT, POD) and photosynthetic pigments (Chl a, Chl b, Total Chl, Car), suggesting that severe growth inhibition in susceptible lines triggers reactive enzymatic and pigment modulation under water deficit. To complement our assessment, we also calculated the Stress Tolerance Index (STI), which captures trait stability under both stress and non-stress conditions and therefore identifies which genotypes are the best overall performers (**Supplemental Figure 1a**). Hierarchical clustering grouped the 31 genotypes into distinct performance clusters based on their overall multi-trait performance under drought. Several Mexican landraces (Mérida, Comala, Guaco, and Xicotepec) exhibited high positive STI scores across the entire biomass trait block (RFW, RDW, PH, SD, PFW, LA, PDW), reflecting high productivity under control conditions coupled with stress resilience under water deficit, suggesting these genotypes may be valuable donors for drought tolerance in tomato. Conversely, the wild species *S. pimpinellifolium*, the F1 hybrid and the heirlooms Amana, Thessaloniki, Amish gold, and our reference M82 displayed higher STI values for physiological traits, showing better performance for LRWC and photosynthetic pigments.

Comparing the results from SSI and STI indexes Stress Susceptibility Index (SSI) and Stress Tolerance Index (STI) revealed contrasting physiological strategies in our population (**Figure 2a and Supplemental Figure 1a**). While top-performing commercial cultivars such as Mérida, Guaco, and Comala maintained high overall productivity (high STI) and low relative biomass loss (low SSI), other lines exhibited distinct trade-offs between growth maintenance and physiological stability, suggesting these genotypes exhibit different molecular responses underlying their physiological differences. To better understand these differences, we mapped the genotypes onto a 2D biplot contrasting Biomass STI against Biomass SSI, which established four distinct functional categories (**Supplemental Figure 1b**). We found the Mexican landraces Guaco, Comala, and Xicotepec, and the heirlooms Prudens Purple and Mortgage lifter were the best performers as they exhibit high STI (better biomass potential) and low SSI (high stability under drought conditions), along with other lines that clustered in Group A. Group B consisted of vigorous lines that suffered significant growth penalties under drought. Notably, we found a cluster of genotypes (Group C), including the CRISPR/Cas9-generated mutants *spro-4* and *slcle9* that maintain low susceptibility to drought despite possessing lower biomass than lines from Group A under non-stress conditions. These mutants were not clustered near their parental background M82 (Group D), confirming they exhibit enhanced drought tolerance. To further complement our analysis, we performed a Principal Component Analysis (PCA) for both control and water deficit conditions (**Supplemental Figure 1c-d**). Our PCA confirmed the phenotypic diversity present in our collection for all 18 scored traits. Under normal conditions, we observed a cluster of lines with larger biomass and higher levels of chlorophyll and carotenoids (Mérida, Rehovot 13, Comala, Mortgage Lifter, Globonnie and Hotset). The most prominent feature of PCA under water deficit was the separation between biochemical/physiological and morphological traits (**Supplemental Figure 1c-d**). The vectors for Carotenoids, Chlorophyll and stress-related enzymes were generally in the opposite direction of biomass traits (PH, SD, LA, fresh and dry weights). These results suggest that plants that are unable to cope with water deficit exhibit increased oxidative stress, carotenoid accumulation and enhanced cell death as evidenced by increased ion leakage. Prudens Purple, Break O’Day, Mérida and Rehovot 13 exhibited higher shoot to root biomass allocation under water deficit conditions (**Supplemental Figure 1c-d**). This analysis also confirmed the CRISPR/Cas9 mutants *slcle9*, *spro-4* and *spro-5* were not clustered together or near their parental genotype M82 in the PCA under normal or drought conditions, indicating these mutants exhibit morphological and physiological differences likely caused by the mutations in genes regulating plant architecture and flowering time.

### The tomato *slcle9* mutant shows increased tolerance to drought stress

Our previous screening found that mutants affected in flowering time and plant architecture exhibited different responses to drought compared to our reference M82 (**Figure 1b-i, Supplemental Figure 1, Figure 2** and **Supplemental Tables 2-7**). Intriguingly, *SlCLE9* was previously found to regulate shoot meristems via transcriptional upregulation upon genetic perturbations of its closest paralog, *SlCLV3*. In our screening, the knockout promoter allele of *SlCLV3* (*slclv3pro*) (Rodríguez-Leal et al. 2017) exhibited poor performance in drought conditions, while the mutant *slcle9* exhibited more stability under drought compared to *slclv3pro* and M82. To further characterize the *slcle9* mutant, we performed three additional experiments comparing the effects of full irrigation (control) and 50% water deficit vs complete water withholding (drought treatment) for 7-11 days, as well as *in vitro* osmotic stress induced by PEG treatment (**Figure 2b-d** and **Figure 3**). As with our previous experiment with 40% water deficit, we observed that *slcle9* exhibited better biomass retention under drought compared to M82 (**Figure 2b-d**).

**Figure 3.**
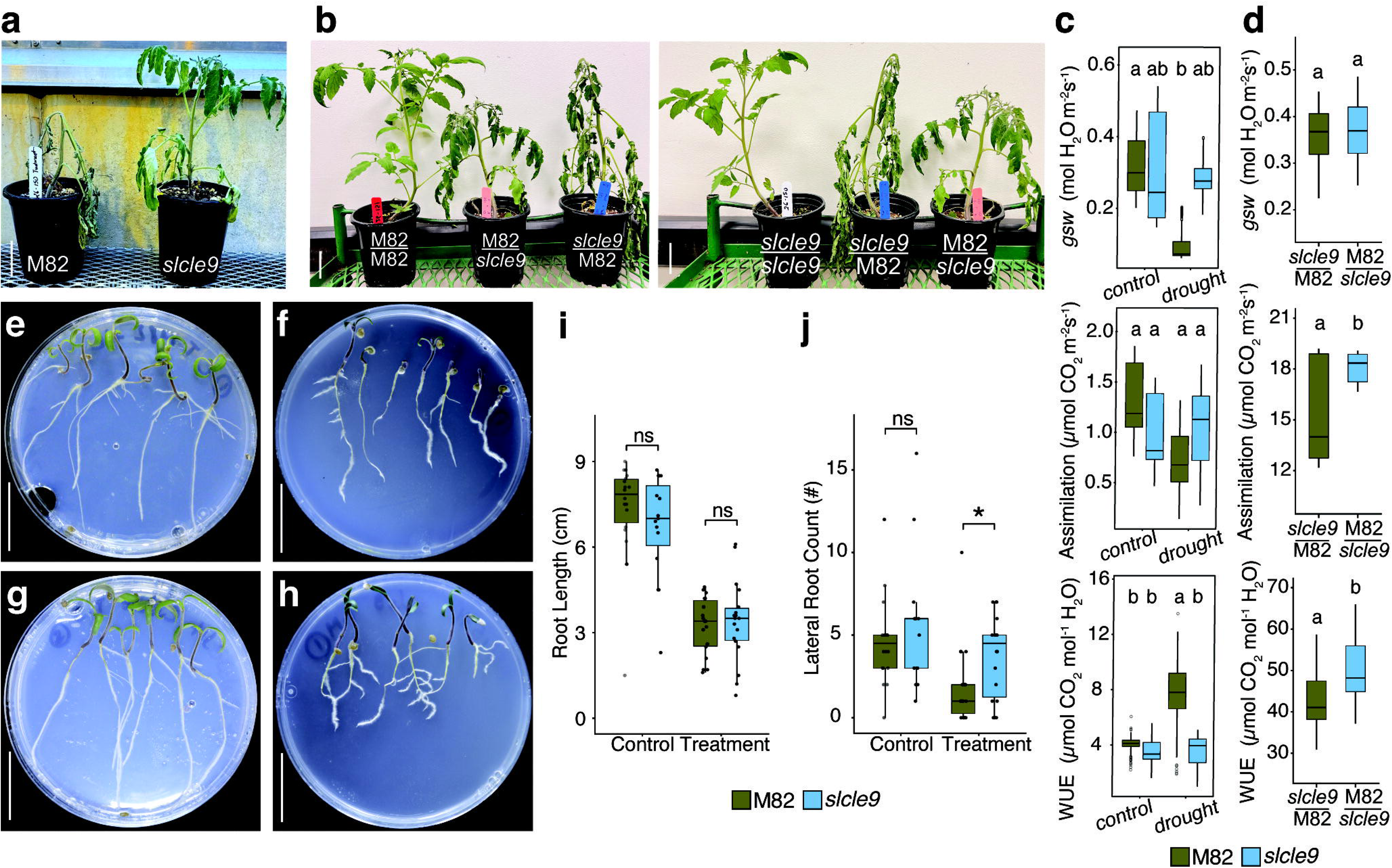
*SlCLE9* is a novel gene involved in drought tolerance. **a**, Representative pictures of M82 (left) and *slcle9* (right) plants after 11 days of complete water withdrawal. **b**, Representative pictures of grafted plants from well-watered self-grafted plants and hetero-grafted plants at 11 days after complete water withholding, demonstrating scion- and rootstock-specific contributions to drought tolerance. Labels indicate scion/rootstock configurations. **c**-**d**, Gas exchange parameters including stomatal conductance (*gsw*), net CO_2_assimilation rate (A), and intrinsic water-use efficiency (WUE) comparing M82 and *slcle9* under control (full irrigation) and drought conditions (11 days after water withdrawal). M82, green; *slcle9*, blue. Different lower-case letters denote statistically significant group differences (*p* < 0.05). **e-h**, Representative pictures of plates containing seedlings grown in media containing MS salts (control) and MS salts plus PEG (treatment). **e, g**, M82 and *slcle9* grown in control plates, respectively. **f, h**, M82 and *slcle9* grown in treatment plates, respectively. **i, j**, Quantitative assessment of root growth comparing M82 (green) and *slcle9* (blue) under control and PEG-induced osmotic stress conditions. Boxplots represent the median, interquartile range, and individual data points. Statistical significance between genotypes denoted as ns = not significant, \**p* < 0.05). Scale bars in **a-b** and **e-h**, 5 cm.

As we observed higher root biomass in the mutant in our 50% water deficit, we reasoned that *slcle9*-mediated drought tolerance may be associated with increased water use efficiency, either through a change in the *slcle9* root architecture or by alterations in leaf stomatal conductance and transpiration. To address this, we tested if *slcle9* could confer drought tolerance by grafting. We grafted *slcle9* as rootstock or scion to its parental genotype M82 and subjected these plants to full irrigation or complete water withholding. Remarkably, both M82 and grafted plants with M82 as rootstocks showed almost complete wilting after 11 days of water withholding, *slcle9* mutants and grafted plants with *slcle9* rootstocks only showed moderate signs of wilting (**Figure 3a-b**). To complement our morphological measurements, we also analyzed stomatal conductance, carbon assimilation and water use efficiency in M82, *slcle9* and grafted plants (**Figure 3c-d**). At 11 days after complete water withholding, *slcle9* exhibited comparable net photosynthesis and stomatal conductance to controls under full irrigation, whereas M82 exhibited reduced assimilation and stomatal conductance (**Figure 3c**). Notably, we observed that scions grafted onto *slcle9* exhibited a trend of lower stomatal conductance but had significantly higher carbon assimilation and water use efficiency at 4 days after complete water withdrawal (**Figure 3d**). These results suggest *SlCLE9* has a novel role in drought tolerance, likely through modifications of root architecture and stomatal conductance under drought stress.

Because of the differences observed between M82 and *slcle9* rootstocks, we decided to study root responses to osmotic stress as a proxy to evaluate drought tolerance by performing *in vitro* assays to induce osmotic stress using PEG (Verslues et al. 2006). Seedlings of M82 and *slcle9* were first germinated in plates containing MS solid media and when roots were emerging (∼5 mm), we transferred them to new plates without PEG (control) and with PEG to achieve a water potential of -0.5 MPa (See **Methods**). After 7 days of growth under controlled chamber conditions, we quantified root lengths and number of lateral roots in both control and treatment plates. While both genotypes showed similar root length and lateral roots under control conditions, *slcle9* exhibited twice the number of lateral roots compared to M82 in the treatment plates (**Figure 3e-j**), suggesting *SlCLE9* has a novel function in controlling root architecture. To better understand the role of *SlCLE9* in root development, we analyzed its expression in M82 and *slcle9* mutants at Day 0 and Day 11 after complete water withholding through quantitative reverse-transcription PCR (qPCR). We observed that *SlCLE9* was upregulated in leaves and roots from M82 and *slcle9* plants at Day 11 after complete water withholding (**Figure 4a**). To further determine the molecular profile of the drought response in *slcle9*, we evaluated 17 genes previously associated with drought responses in tomato and Arabidopsis (Upadhyay et al. 2021). Under non-stressed control conditions, the baseline gene expression in *slcle9* was largely indistinguishable from M82 in both leaves and roots (**Figure 4a**). However, while drought induced severe suppression of light-harvesting complex genes (*Lhcb1*, *Lhcb2*, *Lhcb5*, and *Lhcb6*) in M82 plants, the *slcle9* mutant leaves exhibited attenuated suppression of these photosystem markers under drought, particularly maintaining higher relative expression of *Lhcb1* and *Lhcb2*. Furthermore, the ABA transcriptional regulator *SlDREB2* showed stronger drought-induced upregulation in *slcle9* leaves (log2 FC > 5.0, *p* < 0.001) and pre-elevated baseline levels in roots compared to M82. Similarly, the ABA- and ROS-responsive transcription factor *JUB1* showed stronger drought induction in *slcle9* roots (*p* < 0.001) relative to M82 (*p* < 0.01), while the metabolic marker *ICDH* reached significant upregulation specifically in *slcle9* leaves (*p* < 0.05). These changes were accompanied by heightened expression of ABA biosynthesis genes (*NCED1* and *NCED2*), the stress marker *RD29*, the organic acid metabolic gene *PEPC*, and the lipid/ethylene-related marker *SlaccD* in *slcle9* under drought. Together, these findings demonstrate that *SlCLE9* acts as a crucial regulatory check during water deficit, with its loss leading to attenuated photosynthetic activity and activation of transcriptional, hormonal, and metabolic genes particularly associated with ABA.

**Figure 4.**
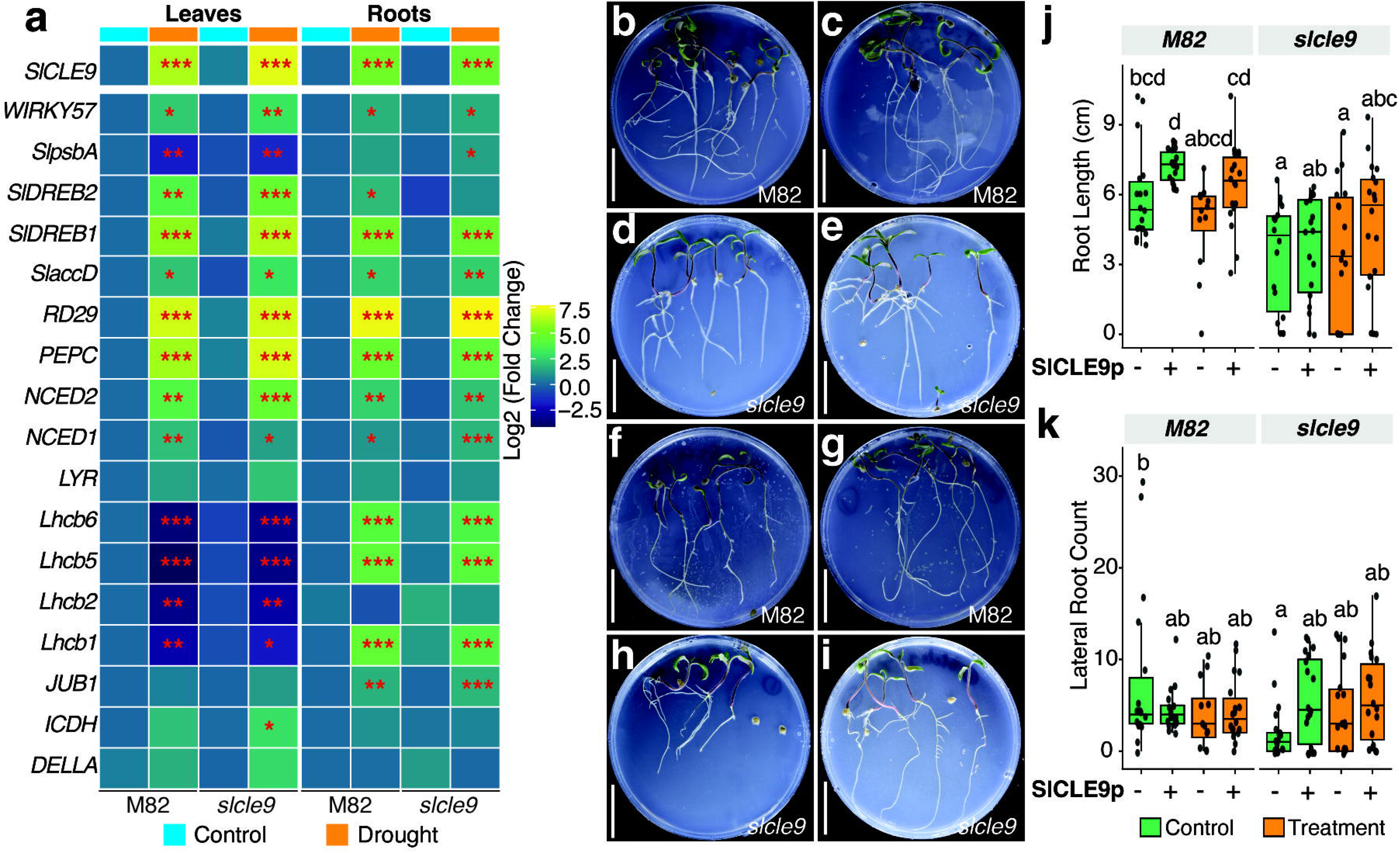
*SlCLE9* is a regulator or root architecture. **a**, Gene expression analysis comparing M82 and *slcle9* under normal (full irrigation) and drought conditions (11 days after complete water withdrawal) using a panel of 18 marker genes including *SlCLE9*. *SlCLE9* upregulation is observed in leaves and roots of both genotypes under drought conditions. Upregulation of ABA-related genes is observed in *slcle9* leaves and roots under stress conditions. Statistical significance is based on ANOVA linear model and p-values are denoted by * (p < 0.05), ** (p < 0.01), and *** (p < 0.001). **b-i**, M82 and *slcle9* seedlings were germinated in media supplemented with PEG and SlCLE9p. **b, d**, M82 and *slcle9* controls (only MS). **c, e**, M82 and *slcle9* controls supplemented with SlCLE9p. **f, h**, M82 and *slcle9* treatment (PEG). **g, i**, M82 and *slcle9* treatment supplemented with SlCLE9p. **j-k**, quantitative data on root length and number of lateral roots from panels **b-i**. Statistical significance values indicated in each plot as group letters from *post-hoc* analysis. Scale bars, **b-i**, 5 cm.

To further determine whether the *SlCLE9* directly regulates root development, we evaluated primary root length and lateral root density in wild-type M82, and *slcle9* seedlings subjected to PEG-induced osmotic stress (-0.5 MPa) with or without an exogenous synthetic SlCLE9 peptide (SlCLE9p) (**Figure 4b-k**). In M82, treatment with SlCLE9p significantly promoted primary root elongation under control conditions (*p* < 0.05), and a similar trend toward enhanced root length was maintained under osmotic stress (**Figure 4b, d, j**). Notably, exogenous application of SlCLE9p restored primary root length in *slcle9* seedlings compared to M82 under both control and osmotic stress conditions, indicating that exogenous SlCLE9 peptide is sufficient to complement root architecture changes in *slcle9* plants under osmotic stress. Similarly, SlCLE9p supplementation restored lateral root initiation in *slcle9* seedlings back to wild-type levels regardless of PEG treatment (**Figure 4k**). Taken together, these results demonstrate that *SlCLE9* is a novel regulator of root development and a modulator of the stress response under drought in the vegetable crop tomato.

## Discussion

Due to more global recurrent and dramatic weather events such as drought, we expect crops will experience yield and quality reductions, effectively threatening food, feed and fuel security. Besides the implementation of management strategies to cope with abiotic stresses, the most efficacious approach to overcome this challenge is by breeding for stress-resilient crops. Tomato is one of the top vegetable crops produced worldwide. Despite its cosmopolitan distribution, it is still very sensitive to drought stress. Previous studies have characterized different cultivars as potential donors for drought-tolerance traits, suggesting tomato has a wide genetic base for breeding stress resilience. Our study is the first showing the response to water deficit in an extended set of commercially relevant heirloom varieties in the Mid-Atlantic US, as well the first characterizing several CRISPR/Cas9 mutants affected in plant architecture and flowering time for their response to drought stress.

At 40% water deficit conditions we observed a significant reduction of biomass (**Figure 1**). Similar negative effects on growth have been reported previously, supporting our experimental design and results (Klunklin and Savage 2017; Parveen et al. 2019; Saidi et al. 2020). Moreover, we observed a broad range of variation across the different traits measured, suggesting our collection harbors wide genetic diversity with potential for improving drought tolerance (**Supplemental Figure 1a-b**). Interestingly, whereas some genotypes such as the Mexican landraces Mérida, Guaco and Xicotepec maintained better biomass under water deficit, others such as the hybrid Mountain Pride F1 and the wild species *S. pimpinellifollium* mounted a stronger physiological response by retaining photosynthetic pigments and carotenoids. Notably, the CRISPR/Cas9 mutants *spro-4* and *slcle9* also showed better root biomass retention (23% and 19% root dry weight reduction, respectively) compared to their reference background M82 (39%, **Supplemental Table 8**), suggesting these mutants previously described to regulate inflorescence architecture and meristem homeostasis have a novel role in drought responses in tomato. Collectively, the negative effect of water deficit was not uniform across the 31 genotypes, with some exhibiting stronger negative effects on growth upon water deficit.

Our study shows that Mexican landraces that come from hot, arid or humid environments from Mexico are potential donors for genes conferring resistance to drought (**Figure 1** and **Supplemental Figure 1**). Future genetic studies with these landraces may uncover novel alleles conferring resistance against these abiotic factors, which are known to exacerbate the effects of drought (Khan et al. 2024; Sellami and Kooli 2026). It is important to acknowledge that our research was focused primarily on assessments of drought tolerance at vegetative stages. Because of the polygenic and *Genotype x Environment* (GxE) interactions that regulate a plant’s response to drought, future evaluations under field conditions or long-term water deficit treatments in reproductive stages will be needed to further confirm the breeding value of these materials.

Notably, we found the heirloom cultivars Prudens purple, Eva purple ball, Mortgage lifter and Cherokee purple exhibited promising tolerance to water deficit at vegetative stages, either by maintaining better leaf water content, higher enzymatic activities or higher biomass. Heirlooms are open-pollinated varieties that have not been subjected to systematic breeding in several decades. Despite this, local communities have made selections based on fruit qualities and characteristics such as flavor profile, color, aroma, fruit size and shape. Due to the accelerated trend of increased temperatures over the last few decades caused by human-induced climate change (Rebecca Lindsey and LuAnn Dahlman), it is likely that local or regional growers were indirectly selecting more stress-resilient heirlooms. Therefore, the genetic variability present in heirlooms may be of potential use for discovery and genetic characterization of novel genes or variants associated with drought tolerance using conventional mapping and genomic approaches, which will allow the development of improved local varieties accessible to US markets, especially for urban agriculture.

Our screening was unique in that we included several previously published CRISPR/Cas9 mutants previously known to regulate shoot architecture, plant growth and flowering time (Xu et al. 2015; Rodríguez-Leal et al. 2017; Soyk et al. 2017; Rodriguez-Leal et al. 2019). Notably, we discovered that the mutant *slcle9* exhibited improved response to water deficit compared to its parental genotype M82. *SlCLE9* encodes for the closest paralog of the shoot apical meristem stem-cell regulator *SlCLAVATA3* (*SlCLV3*) (Xu et al. 2015). Previously, we demonstrated that *slcle9* mutants exhibited a very modest increase in locule number that was dramatically exacerbated in the *slclv3* mutant background, showing that *SlCLE9* acts as a compensator regulating stem cell proliferation (Rodriguez-Leal et al. 2019). However, no other phenotypes have been associated with mutations in *SlCLE9*. Here, we found that while *slclv3* mutants were more susceptible to drought, *slcle9* showed enhanced drought tolerance (**Figure 2** and **Supplemental Figure 1**). This type of “unequal redundancy” suggests that *SlCLV3* may have evolved primarily to regulate stem cell proliferation, while *SlCLE9* may have evolved novel functions in plant development. As some Solanaceous crops such as pepper have lost *SlCLE9* paralog, while others (*Physalis spp*) have retained both paralogs (Rodriguez-Leal et al. 2019; Kwon et al. 2022), further studies may reveal the evolutionary path that led to divergence in *SlCLE9* function and the mechanisms of paralog retention based on neo- or subfunctionalization.

Our research is the first demonstrating the role of *SlCLE9* in drought tolerance in the vegetable crop tomato. Although the *slcle9* mutant allele carries a deletion in the 3’ end of the transcript, effectively disrupting the functional CLE (dodecapeptide) domain, the mutant exhibited a strong transcriptional response in mutant backgrounds carrying different alleles of *SlCLV3* (Xu et al. 2015; Rodriguez-Leal et al. 2019; Wang et al. 2021; Aguirre et al. 2023), suggesting that *SlCLE9* may have regulatory regions with *cis*-elements that may respond to genetic or environmental perturbations. Our expression analysis confirms *SlCLE9* is responding transcriptionally to drought stress (**Figure 4a**), suggesting the presence of motifs in its *cis*-regulatory region associated with transcription factors that regulate drought in tomato. The *CLAVATA3/EMBRYO SURROUNDING REGION* (*CLE*) gene family is diverse and composed of dozens of members across different plant species (Rodriguez-Leal et al. 2019; Carbonnel et al. 2022; Gentile et al. 2025). Many CLE peptides have been shown to regulate responses to drought, heat or salt stress in the model species *Arabidopsis thaliana*. (Cao et al., 2025). Compared to recent reports showing the role of CLE peptides in tomato and Arabidopsis (Endo and Fukuda 2024; Wang et al. 2026) our study shows that *SlCLE9* is likely modulating both root and shoot responses to drought stress. Indeed, we observed the upregulation of drought-related genes, particularly associated with Abscisic Acid (ABA) signaling, suggesting *SlCLE9* acts as a negative feedback regulator or systemic signal modulating the threshold of ABA and stress gene activation. This is supported by our physiological measurements comparing *M82* and *slcle9* and by the results of our *in vitro* peptide treatment. When seedlings are exposed to the *SlCLE9* peptide in the growth media, both M82 and *slcle9* exhibited larger primary roots, even under drought/osmotic stress (**Figure 4b-k**). Recent genetics studies suggest that CLV1 and related BAM receptors may be involved in drought responses in Arabidopsis (Bashyal et al. 2023), and we previously demonstrated the peptide SlCLE9 is likely perceived by the receptor CLV1 (Rodriguez-Leal et al. 2019). Thus, it seems plausible that *SlCLE9* may be functioning as a modulator of the stress response, signaling through the *CLV1* or other related receptors, as previously shown in Arabidopsis (Smith et al. 2025), including for the response to nutrient deficiency, which also triggered overexpression of several Arabidopsis *CLE* genes (Araya et al. 2014). Whether this role is directly associated with signaling from shoot to root, particularly involving the shoot apical meristem, or by a novel role of *SlCLE9* in root development and physiology is still unknown. Remarkably, we demonstrated that when *slcle9* was used as rootstock, the scions of M82 were more resilient to drought stress. Rootstock breeding programs are limited to relatively few donor lines from wild species (Lee et al. 2023). Our research offers a new venue for breeding for drought tolerance via mutagenesis of *SlCLE9*. Our results demonstrate a novel role for *SlCLE9* as a modulator of the response to drought stress in the vegetable crop tomato with potential for improvement in rootstock breeding and urban agriculture programs.

## Supporting information

Supp Datafile 1

## Acknowledgements

We thank all members of the Tomato Lab for their support on this research project. We thank Sydney Wallace, Meghan Fisher, and Christian Mckay for their assistance in greenhouse plant care. We thank Dr. Zach Lippman (CSHL) for sharing mutant seeds of *slcle9* and other mutants used in this study. We thank the TGRC for generously providing stocks of the different tomato cultivars used in this study. We thank Justin Liu, Guthrie Specht and Gabriel Villareal for their help preparing reagents for the *in vitro* experiments. We thank Dr. John Erwin for sharing Department equipment for our physiology experiments. This research was supported by grants from MAES (MD-PSLA-243084), the California Tomato Research Institute (CTRI), startup funds from UMD-PSLA, and a postdoctoral fellowship to M.E. from Tubitak (TUBİTAK 2219/ 1059B192402353). RKU research at Bowie State University was supported by new lab space provided by the Department of Natural Sciences and funding support from a faculty development grant (GR23001023).

## Author contributions

M.E., S.O., H.L., R.W, D.R.-L, A.A, J.O. and C.A.S. performed drought experiments; A.A, S.C.G and R.K.U. performed expression analysis; M.E. and D.R.-L. conceived the project, selected the cultivars for the screening, supervised all experiments and performed greenhouse care. M.E. and D.R.-L. wrote and edited the figures and manuscript. All authors read and approved the manuscript.

---

**Supplemental Figure 1.**
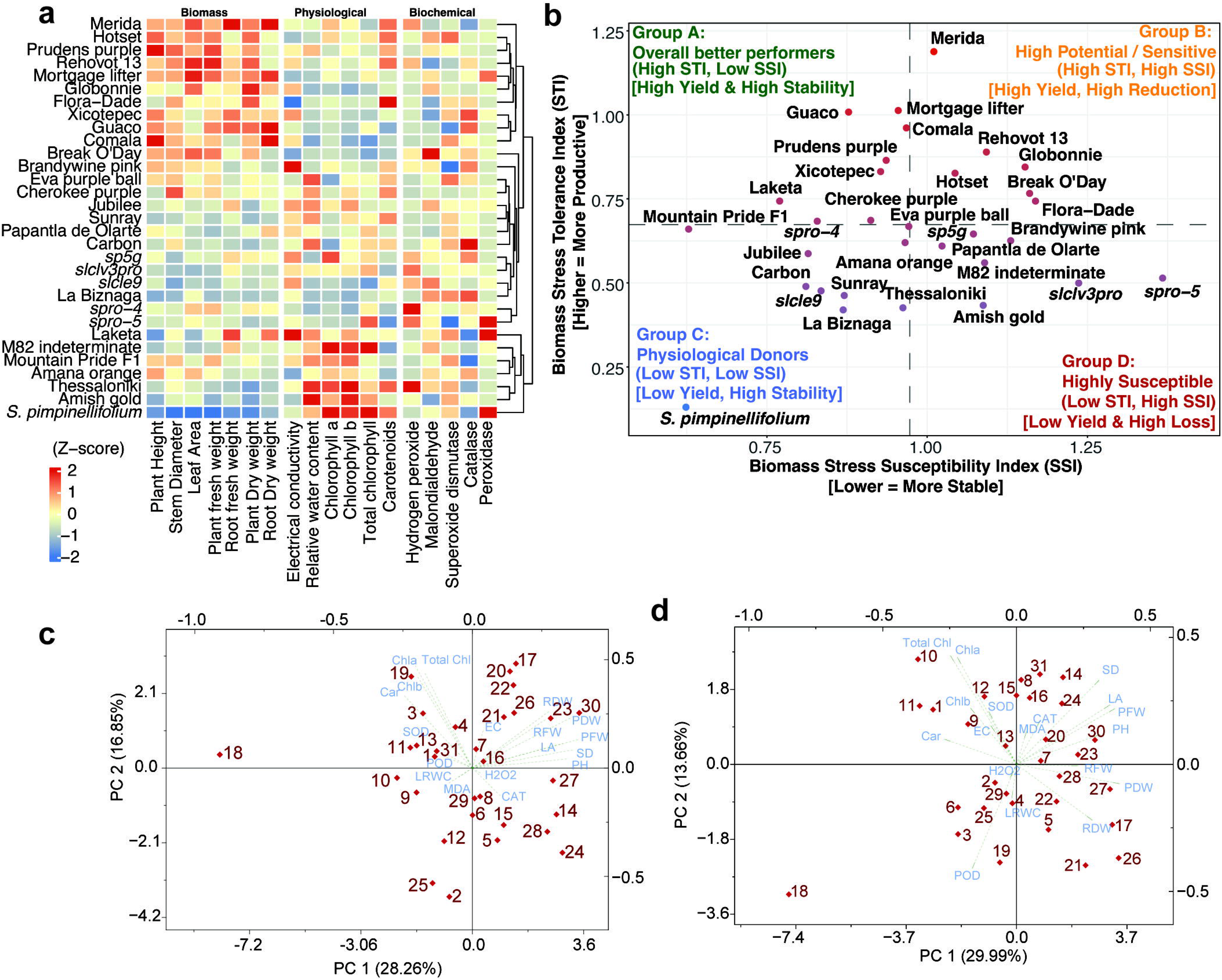
Integrated stress indices and multivariate profiling of the population response to water deficit. **a**, Heatmap of column-standardized (Z-score) Stress Tolerance Index (STI) values across 18 phenotypic traits categorized into Biomass, Physiological, and Biochemical functional groups. Genotypes (rows) are hierarchically clustered according to multi-trait STI similarity, while traits within functional blocks maintain standardized grouping. Color scale denotes relative stress tolerance potential, ranging from low (blue, negative Z-scores) to high (red, positive Z-scores). **b**, Four-quadrant biplot classifying the 31 tomato genotypes used in this study based on Biomass Stress Tolerance Index (STI; y-axis, higher indicates greater productivity) versus Biomass Stress Susceptibility Index (SSI; x-axis, lower indicates greater stability). Quadrants categorize lines into Group A (Gold Standard: high STI, low SSI; high yield and stability), Group B (High Potential/Sensitive: high STI, high SSI; high yield but high reduction), Group C (Physiological Donors: low STI, low SSI; low yield but high stability), and Group D (Highly Susceptible: low STI, high SSI; low yield and severe loss). **c-d**, Principal Component Analysis (PCA) biplots illustrating trait vectors and individual distributions of the 31 tomato genotypes grown under well-watered control conditions (**c**) and under water deficit conditions (**d**). In both condition-specific PCAs, blue vectors indicate trait loadings, and numbered points represent individual tomato genotypes. PC1 and PC2 account for 28.26% and 16.85% of total variance under control conditions (**c**), and 29.99% and 13.66% under water deficit (**d**), respectively.

