## Supplementary material for "Morphological and biochemical evaluation of tolerance to water deficit identifies *SlCLE9* as a novel regulator of drought stress": Supp Datafile 1

#### The GLM Procedure

#### Class Level Information

| Class | Levels | Values |
| --- | --- | --- |
| Genotype | 31 | 1 2 3 4 5 6 7 8 9 10 11 12 13 14 15 16 17 18 19 20 21 22 23 24 25 26 27 28 29 30 31 |
| FC____ | 2 | 60 100 |

Number of observations 186

14:00 Thursday, February 18, 2026 2

#### The GLM Procedure

Dependent Variable: PH\_\_cm\_ PH (cm)

| Source | DF | Sum of Squares | Mean Square | F Value | Pr > F |
| --- | --- | --- | --- | --- | --- |
| Model | 61 | 4092.302628 | 67.086928 | 72.25 | <.0001 |
| Error | 124 | 115.138667 | 0.928538 |  |  |
| Corrected Total | 185 | 4207.441295 |  |  |  |

|  |  |  |  |
| --- | --- | --- | --- |
| R-Square | Coeff Var | Root MSE | PH__cm_ Mean |
| 0.972635 | 3.884584 | 0.963607 | 24.80591 |

| Source | DF | Type III SS | Mean Square | F Value | Pr > F |
| --- | --- | --- | --- | --- | --- |
| Genotype | 30 | 2009.584728 | 66.986158 | 72.14 | <.0001 |
| FC____ | 1 | 1702.681815 | 1702.681815 | 1833.72 | <.0001 |
| Genotype*FC____ | 30 | 380.036085 | 12.667869 | 13.64 | <.0001 |

14:00 Thursday, February 18, 2026 3

#### The GLM Procedure

Dependent Variable: SD\_\_cm\_ SD (cm)

| Source | DF | Sum of Squares | Mean Square | F Value | Pr > F |
| --- | --- | --- | --- | --- | --- |
| Model | 61 | 20.46380914 | 0.33547228 | 82.08 | <.0001 |
| Error | 124 | 0.50680000 | 0.00408710 |  |  |
| Corrected Total | 185 | 20.97060914 |  |  |  |

|  |  |  |  |
| --- | --- | --- | --- |
| R-Square | Coeff Var | Root MSE | SD__cm_ Mean |
| 0.975833 | 3.117494 | 0.063930 | 2.050699 |

| Source | DF | Type III SS | Mean Square | F Value | Pr > F |
| --- | --- | --- | --- | --- | --- |
| Genotype | 30 | 11.45995914 | 0.38199864 | 93.46 | <.0001 |
| FC____ | 1 | 7.76752742 | 7.76752742 | 1900.50 | <.0001 |
| Genotype*FC____ | 30 | 1.23632258 | 0.04121075 | 10.08 | <.0001 |

14:00 Thursday, February 18, 2026 4

#### The GLM Procedure

Dependent Variable: LA\_\_cm2\_ LA (cm2)

| Source | DF | Sum of Squares | Mean Square | F Value | Pr > F |
| --- | --- | --- | --- | --- | --- |
| Model | 61 | 2154278.713 | 35316.044 | 131.62 | <.0001 |
| Error | 124 | 33272.490 | 268.327 |  |  |
| Corrected Total | 185 | 2187551.203 |  |  |  |

|  |  |  |  |
| --- | --- | --- | --- |
| R-Square | Coeff Var | Root MSE | LA_cm2_ Mean |
| 0.984790 | 4.907390 | 16.38068 | 333.7961 |

|  |  |  |  |  |  |
| --- | --- | --- | --- | --- | --- |
| Source | DF | Type III SS | Mean Square | F Value | Pr > F |
| Genotype | 30 | 730122.279 | 24337.409 | 90.70 | <.0001 |
| FC_____ | 1 | 1212971.594 | 1212971.594 | 4520.51 | <.0001 |
| Genotype*FC_____ | 30 | 211184.840 | 7039.495 | 26.23 | <.0001 |

14:00 Thursday, February 18, 2026 5

The GLM Procedure

Dependent Variable: PFW\_\_g\_ PFW (g)

|  |  |  |  |  |  |
| --- | --- | --- | --- | --- | --- |
| Source | DF | Sum of Squares | Mean Square | F Value | Pr > F |
| Model | 61 | 5103.891398 | 83.670351 | 181.83 | <.0001 |
| Error | 124 | 57.060000 | 0.460161 |  |  |
| Corrected Total | 185 | 5160.951398 |  |  |  |

|  |  |  |  |
| --- | --- | --- | --- |
| R-Square | Coeff Var | Root MSE | PFW__g_ Mean |
| 0.988944 | 4.153995 | 0.678352 | 16.33011 |

|  |  |  |  |  |  |
| --- | --- | --- | --- | --- | --- |
| Source | DF | Type III SS | Mean Square | F Value | Pr > F |
| Genotype | 30 | 2045.634731 | 68.187824 | 148.18 | <.0001 |
| FC_____ | 1 | 2641.940860 | 2641.940860 | 5741.34 | <.0001 |
| Genotype*FC_____ | 30 | 416.315806 | 13.877194 | 30.16 | <.0001 |

14:00 Thursday, February 18, 2026 6

The GLM Procedure

Dependent Variable: RFW\_\_g\_ RFW (g)

|  |  |  |  |  |  |
| --- | --- | --- | --- | --- | --- |
| Source | DF | Sum of Squares | Mean Square | F Value | Pr > F |
| Model | 61 | 973.811606 | 15.964125 | 40.06 | <.0001 |
| Error | 124 | 49.414400 | 0.398503 |  |  |
| Corrected Total | 185 | 1023.226006 |  |  |  |

|  |  |  |  |
| --- | --- | --- | --- |
| R-Square | Coeff Var | Root MSE | RFW__g_ Mean |
| 0.951707 | 10.95098 | 0.631271 | 5.764516 |

|  |  |  |  |  |  |
| --- | --- | --- | --- | --- | --- |
| Source | DF | Type III SS | Mean Square | F Value | Pr > F |
| Genotype | 30 | 466.5270731 | 15.5509024 | 39.02 | <.0001 |
| FC_____ | 1 | 396.8865054 | 396.8865054 | 995.94 | <.0001 |
| Genotype*FC_____ | 30 | 110.3980280 | 3.6799343 | 9.23 | <.0001 |

14:00 Thursday, February 18, 2026 7

The GLM Procedure

Dependent Variable: PDW\_\_g\_ PDW (g)

|  |  |  |  |  |  |
| --- | --- | --- | --- | --- | --- |
| Source | DF | Sum of Squares | Mean Square | F Value | Pr > F |
| Model | 61 | 64.90258280 | 1.06397677 | 115.22 | <.0001 |
| Error | 124 | 1.14506667 | 0.00923441 |  |  |
| Corrected Total | 185 | 66.04764946 |  |  |  |

|  |  |  |  |
| --- | --- | --- | --- |
| R-Square | Coeff Var | Root MSE | PDW__g_ Mean |
| 0.982663 | 6.954796 | 0.096096 | 1.381720 |

| Source | DF | Type III SS | Mean Square | F Value | Pr > F |
| --- | --- | --- | --- | --- | --- |
| Genotype | 30 | 28.97894946 | 0.96596498 | 104.60 | <.0001 |
| FC_____ | 1 | 31.33223441 | 31.33223441 | 3392.99 | <.0001 |
| Genotype*FC_____ | 30 | 4.59139892 | 0.15304663 | 16.57 | <.0001 |

14:00 Thursday, February 18, 2026 8

###### The GLM Procedure

Dependent Variable: RDW\_\_g\_ RDW (g)

| Source | DF | Sum of Squares | Mean Square | F Value | Pr > F |
| --- | --- | --- | --- | --- | --- |
| Model | 61 | 5.75112312 | 0.09428071 | 60.06 | <.0001 |
| Error | 124 | 0.19466667 | 0.00156989 |  |  |
| Corrected Total | 185 | 5.94578978 |  |  |  |

R-Square Coeff Var Root MSE RDW\_\_g\_ Mean
0.967260 11.19840 0.039622 0.353817

| Source | DF | Type III SS | Mean Square | F Value | Pr > F |
| --- | --- | --- | --- | --- | --- |
| Genotype | 30 | 3.08410645 | 0.10280355 | 65.48 | <.0001 |
| FC_____ | 1 | 2.08862419 | 2.08862419 | 1330.42 | <.0001 |
| Genotype*FC_____ | 30 | 0.57839247 | 0.01927975 | 12.28 | <.0001 |

14:00 Thursday, February 18, 2026 9

###### The GLM Procedure

Dependent Variable: EC\_\_\_\_\_ EC ( )

| Source | DF | Sum of Squares | Mean Square | F Value | Pr > F |
| --- | --- | --- | --- | --- | --- |
| Model | 61 | 38939.47021 | 638.35197 | 49.89 | <.0001 |
| Error | 124 | 1586.74705 | 12.79635 |  |  |
| Corrected Total | 185 | 40526.21726 |  |  |  |

R-Square Coeff Var Root MSE EC\_\_\_\_\_ Mean
0.960846 11.25495 3.577198 31.78333

| Source | DF | Type III SS | Mean Square | F Value | Pr > F |
| --- | --- | --- | --- | --- | --- |
| Genotype | 30 | 11928.68049 | 397.62268 | 31.07 | <.0001 |
| FC_____ | 1 | 17837.52595 | 17837.52595 | 1393.95 | <.0001 |
| Genotype*FC_____ | 30 | 9173.26378 | 305.77546 | 23.90 | <.0001 |

14:00 Thursday, February 18, 2026 10

###### The GLM Procedure

Dependent Variable: LRWC\_\_\_\_\_ LRWC ( )

| Source | DF | Sum of Squares | Mean Square | F Value | Pr > F |
| --- | --- | --- | --- | --- | --- |
| Model | 61 | 6881.31028 | 112.80837 | 3.41 | <.0001 |
| Error | 124 | 4105.92907 | 33.11233 |  |  |
| Corrected Total | 185 | 10987.23934 |  |  |  |

R-Square Coeff Var Root MSE LRWC\_\_\_\_\_ Mean
0.626300 8.631015 5.754332 66.67039

| Source | DF | Type III SS | Mean Square | F Value | Pr > F |
| --- | --- | --- | --- | --- | --- |

|  |  |  |  |  |  |
| --- | --- | --- | --- | --- | --- |
| Genotype | 30 | 2508.848071 | 83.628269 | 2.53 | 0.0002 |
| FC_____ | 1 | 3588.027027 | 3588.027027 | 108.36 | <.0001 |
| Genotype*FC_____ | 30 | 784.435179 | 26.147839 | 0.79 | 0.7700 |

14:00 Thursday, February 18, 2026 11

The GLM Procedure

Dependent Variable: Chla\_\_mg\_g\_1\_ Chla (mg g-1)

| Source | DF | Sum of Squares | Mean Square | F Value | Pr > F |
| --- | --- | --- | --- | --- | --- |
| Model | 61 | 391.5126143 | 6.4182396 | 53.41 | <.0001 |
| Error | 124 | 14.9015924 | 0.1201741 |  |  |
| Corrected Total | 185 | 406.4142067 |  |  |  |

|  |  |  |  |
| --- | --- | --- | --- |
| R-Square | Coeff Var | Root MSE | Chla__mg_g_1_ Mean |
| 0.963334 | 7.730813 | 0.346661 | 4.484152 |

| Source | DF | Type III SS | Mean Square | F Value | Pr > F |
| --- | --- | --- | --- | --- | --- |
| Genotype | 30 | 76.1006190 | 2.5366873 | 21.11 | <.0001 |
| FC_____ | 1 | 254.4535596 | 254.4535596 | 2117.37 | <.0001 |
| Genotype*FC_____ | 30 | 60.9584358 | 2.0319479 | 16.91 | <.0001 |

14:00 Thursday, February 18, 2026 12

The GLM Procedure

Dependent Variable: Chlb\_\_mg\_g\_1\_ Chlb (mg g-1)

| Source | DF | Sum of Squares | Mean Square | F Value | Pr > F |
| --- | --- | --- | --- | --- | --- |
| Model | 61 | 248.6365253 | 4.0760086 | 157.27 | <.0001 |
| Error | 124 | 3.2138336 | 0.0259180 |  |  |
| Corrected Total | 185 | 251.8503589 |  |  |  |

|  |  |  |  |
| --- | --- | --- | --- |
| R-Square | Coeff Var | Root MSE | Chlb__mg_g_1_ Mean |
| 0.987239 | 6.131343 | 0.160991 | 2.625701 |

| Source | DF | Type III SS | Mean Square | F Value | Pr > F |
| --- | --- | --- | --- | --- | --- |
| Genotype | 30 | 64.1417800 | 2.1380593 | 82.49 | <.0001 |
| FC_____ | 1 | 142.9902989 | 142.9902989 | 5517.02 | <.0001 |
| Genotype*FC_____ | 30 | 41.5044464 | 1.3834815 | 53.38 | <.0001 |

14:00 Thursday, February 18, 2026 13

The GLM Procedure

Dependent Variable: Total\_Ch1\_\_mg\_g\_1\_ Total Chl (mg g-1)

| Source | DF | Sum of Squares | Mean Square | F Value | Pr > F |
| --- | --- | --- | --- | --- | --- |
| Model | 61 | 1114.623740 | 18.272520 | 107.17 | <.0001 |
| Error | 124 | 21.142063 | 0.170501 |  |  |
| Corrected Total | 185 | 1135.765803 |  |  |  |

|  |  |  |  |
| --- | --- | --- | --- |
| R-Square | Coeff Var | Root MSE | Total_Chl__mg_g_1_ Mean |
| 0.981385 | 5.808608 | 0.412917 | 7.108709 |

| Source | DF | Type III SS | Mean Square | F Value | Pr > F |
| --- | --- | --- | --- | --- | --- |
| Genotype | 30 | 173.6283347 | 5.7876112 | 33.94 | <.0001 |
| FC_____ | 1 | 778.4899507 | 778.4899507 | 4565.91 | <.0001 |
| Genotype*FC_____ | 30 | 162.5054547 | 5.4168485 | 31.77 | <.0001 |

14:00 Thursday, February 18, 2026 14

#### The GLM Procedure

Dependent Variable: Car\_\_mg\_g\_1\_ Car (mg g-1)

| Source | DF | Sum of Squares | Mean Square | F Value | Pr > F |
| --- | --- | --- | --- | --- | --- |
| Model | 61 | 1332.050130 | 21.836887 | 20.11 | <.0001 |
| Error | 124 | 134.652647 | 1.085908 |  |  |
| Corrected Total | 185 | 1466.702777 |  |  |  |

| R-Square | Coeff Var | Root MSE | Car__mg_g_1_ Mean |
| --- | --- | --- | --- |
| 0.908194 | 8.722959 | 1.042069 | 11.94628 |

| Source | DF | Type III SS | Mean Square | F Value | Pr > F |
| --- | --- | --- | --- | --- | --- |
| Genotype | 30 | 226.6040519 | 7.5534684 | 6.96 | <.0001 |
| FC____ | 1 | 880.6457390 | 880.6457390 | 810.98 | <.0001 |
| Genotype*FC____ | 30 | 224.8003391 | 7.4933446 | 6.90 | <.0001 |

14:00 Thursday, February 18, 2026 15

#### The GLM Procedure

Dependent Variable: H2O2\_\_mmol\_kg\_1\_ H2O2\_(mmol kg-1)

| Source | DF | Sum of Squares | Mean Square | F Value | Pr > F |
| --- | --- | --- | --- | --- | --- |
| Model | 61 | 10742.47646 | 176.10617 | 738.89 | <.0001 |
| Error | 124 | 29.55416 | 0.23834 |  |  |
| Corrected Total | 185 | 10772.03062 |  |  |  |

| R-Square | Coeff Var | Root MSE | H2O2__mmol_kg_1_ Mean |
| --- | --- | --- | --- |
| 0.997256 | 4.836748 | 0.488201 | 10.09357 |

| Source | DF | Type III SS | Mean Square | F Value | Pr > F |
| --- | --- | --- | --- | --- | --- |
| Genotype | 30 | 8349.739744 | 278.324658 | 1167.76 | <.0001 |
| FC____ | 1 | 1392.504753 | 1392.504753 | 5842.51 | <.0001 |
| Genotype*FC____ | 30 | 1000.231965 | 33.341066 | 139.89 | <.0001 |

14:00 Thursday, February 18, 2026 16

#### The GLM Procedure

Dependent Variable: MDA\_\_mmol\_kg\_1\_ MDA\_ (mmol kg-1)

| Source | DF | Sum of Squares | Mean Square | F Value | Pr > F |
| --- | --- | --- | --- | --- | --- |
| Model | 61 | 359.3157452 | 5.8904221 | 173.25 | <.0001 |
| Error | 124 | 4.2158515 | 0.0339988 |  |  |
| Corrected Total | 185 | 363.5315968 |  |  |  |

| R-Square | Coeff Var | Root MSE | MDA__mmol_kg_1_ Mean |
| --- | --- | --- | --- |
| 0.988403 | 4.618943 | 0.184388 | 3.991988 |

| Source | DF | Type III SS | Mean Square | F Value | Pr > F |
| --- | --- | --- | --- | --- | --- |
| Genotype | 30 | 76.4670945 | 2.5489032 | 74.97 | <.0001 |
| FC____ | 1 | 211.0902328 | 211.0902328 | 6208.75 | <.0001 |
| Genotype*FC____ | 30 | 71.7584179 | 2.3919473 | 70.35 | <.0001 |

14:00 Thursday, February 18, 2026 17

#### The GLM Procedure

Dependent Variable: SOD\_\_EU\_g\_1\_Leaf\_ SOD\_(EU g-1 Leaf)

| Source | DF | Sum of Squares | Mean Square | F Value | Pr > F |
| --- | --- | --- | --- | --- | --- |
| Model | 61 | 78141.11499 | 1281.00189 | 87.59 | <.0001 |
| Error | 124 | 1813.48519 | 14.62488 |  |  |
| Corrected Total | 185 | 79954.60017 |  |  |  |

|  |  |  |  |
| --- | --- | --- | --- |
| R-Square | Coeff Var | Root MSE | SOD__EU_g_1_Leaf_ Mean |
| 0.977319 | 1.220521 | 3.824249 | 313.3292 |

| Source | DF | Type III SS | Mean Square | F Value | Pr > F |
| --- | --- | --- | --- | --- | --- |
| Genotype | 30 | 12652.61814 | 421.75394 | 28.84 | <.0001 |
| FC_____ | 1 | 54195.77062 | 54195.77062 | 3705.72 | <.0001 |
| Genotype*FC_____ | 30 | 11292.72624 | 376.42421 | 25.74 | <.0001 |

14:00 Thursday, February 18, 2026 18

###### The GLM Procedure

Dependent Variable: CAT\_\_EU\_g\_1\_Leaf\_ CAT (EU g-1 Leaf)

| Source | DF | Sum of Squares | Mean Square | F Value | Pr > F |
| --- | --- | --- | --- | --- | --- |
| Model | 61 | 192929.8449 | 3162.7843 | 76.30 | <.0001 |
| Error | 124 | 5140.1501 | 41.4528 |  |  |
| Corrected Total | 185 | 198069.9950 |  |  |  |

|  |  |  |  |
| --- | --- | --- | --- |
| R-Square | Coeff Var | Root MSE | CAT__EU_g_1_Leaf_ Mean |
| 0.974049 | 6.335863 | 6.438387 | 101.6181 |

| Source | DF | Type III SS | Mean Square | F Value | Pr > F |
| --- | --- | --- | --- | --- | --- |
| Genotype | 30 | 54834.43293 | 1827.81443 | 44.09 | <.0001 |
| FC_____ | 1 | 98748.11008 | 98748.11008 | 2382.18 | <.0001 |
| Genotype*FC_____ | 30 | 39347.30193 | 1311.57673 | 31.64 | <.0001 |

14:00 Thursday, February 18, 2026 19

###### The GLM Procedure

Dependent Variable: POD\_\_EU\_g\_1\_Leaf\_ POD (EU g-1 Leaf)

| Source | DF | Sum of Squares | Mean Square | F Value | Pr > F |
| --- | --- | --- | --- | --- | --- |
| Model | 61 | 6944405072 | 113842706 | 3417.80 | <.0001 |
| Error | 124 | 4130288 | 33309 |  |  |
| Corrected Total | 185 | 6948535361 |  |  |  |

|  |  |  |  |
| --- | --- | --- | --- |
| R-Square | Coeff Var | Root MSE | POD__EU_g_1_Leaf_ Mean |
| 0.999406 | 4.037548 | 182.5069 | 4520.242 |

| Source | DF | Type III SS | Mean Square | F Value | Pr > F |
| --- | --- | --- | --- | --- | --- |
| Genotype | 30 | 3734361691 | 124478723 | 3737.11 | <.0001 |
| FC_____ | 1 | 1148095326 | 1148095326 | 34468.3 | <.0001 |
| Genotype*FC_____ | 30 | 2061948055 | 68731602 | 2063.47 | <.0001 |

14:00 Thursday, February 18, 2026 20

###### The GLM Procedure

t Tests (LSD) for PH\_\_cm\_

NOTE: This test controls the Type I comparisonwise error rate, not the experimentwise error rate.

Means with the same letter are not significantly different.

| t Grouping |  |  |  | Mean | N | Genotype |
| --- | --- | --- | --- | --- | --- | --- |
| C<br>C<br>C<br>C<br>C<br>C<br>C<br>C<br>C<br>G<br>G<br>G<br>G<br>G<br>G | F | A |  | 31.0967 | 6 | 26 |
|  |  | A |  |  |  |  |
|  |  | A |  | 30.7500 | 6 | 14 |
|  |  | B |  | 29.3050 | 6 | 22 |
|  |  | B |  |  |  |  |
|  |  | B |  | 28.5417 | 6 | 28 |
|  |  | D |  | 28.1117 | 6 | 27 |
|  |  | D |  |  |  |  |
|  |  | D | E | 27.6383 | 6 | 24 |
|  |  | D | E |  |  |  |
|  |  | D | E | 27.5967 | 6 | 8 |
|  |  | D | E |  |  |  |
|  |  | D | E | 27.5700 | 6 | 30 |
|  |  | D | E |  |  |  |
|  |  | D | E | 27.2500 | 6 | 17 |
|  |  | D | E |  |  |  |
| F | D | E | 27.1667 | 6 | 7 |  |
| F | D | E |  |  |  |  |
| F | D | E | 27.2500 | 6 | 17 |  |
| F | D | E |  |  |  |  |
| F | D | E | 27.1667 | 6 | 7 |  |
| F | D | E |  |  |  |  |
| F | H | E | 26.8467 | 6 | 23 |  |
| F | H |  |  |  |  |  |
| F | H |  | 26.4717 | 6 | 31 |  |
| F | H |  |  |  |  |  |
| F | H |  | 26.1383 | 6 | 29 |  |
| F | H |  |  |  |  |  |
| F | H |  | 26.0417 | 6 | 16 |  |
| F | H |  |  |  |  |  |
| F | H |  | 25.8333 | 6 | 21 |  |
| F | H |  |  |  |  |  |
| J<br>J<br>J<br>J<br>J |  | I |  | 24.6800 | 6 | 4 |
|  |  | I |  |  |  |  |
|  |  | I |  | 24.1667 | 6 | 5 |
|  |  | K |  | 23.4300 | 6 | 6 |
|  |  | K |  |  |  |  |
|  |  | K | L | 23.3633 | 6 | 2 |
|  |  | K | L |  |  |  |

14:00 Thursday, February 18, 2026 21

#### The GLM Procedure

#### t Tests (LSD) for PH\_\_cm\_

Means with the same letter are not significantly different.

| t | Grouping | Mean | N | Genotype |
| --- | --- | --- | --- | --- |
|  | K L | 22.9867 | 6 | 13 |
|  | K L |  |  |  |
|  | K L | 22.9583 | 6 | 20 |
|  | K L |  |  |  |
|  | K L | 22.7500 | 6 | 3 |
|  | K L |  |  |  |
|  | K L | 22.6100 | 6 | 12 |
|  | K L |  |  |  |
|  | K L | 22.5567 | 6 | 15 |
|  | L |  |  |  |
|  | M | 22.2767 | 6 | 10 |
|  | M |  |  |  |
|  | M | 21.4300 | 6 | 1 |
|  | M |  |  |  |
|  | M | 21.3750 | 6 | 9 |
|  | N | 20.2500 | 6 | 11 |
|  | N |  |  |  |
|  | N | 20.0417 | 6 | 25 |
|  | O | 18.9167 | 6 | 19 |

The GLM Procedure

t Tests (LSD) for SD\_cm\_

NOTE: This test controls the Type I comparisonwise error rate, not the experimentwise error rate.

Alpha 0.05
Error Degrees of Freedom 124
Error Mean Square 0.004087
Critical Value of t 1.97928
Least Significant Difference 0.0731

Means with the same letter are not significantly different.

| t Grouping |  | Mean | N | Genotype |
| --- | --- | --- | --- | --- |
|  | A | 2.36333 | 6 | 15 |
|  | A |  |  |  |
| B | A | 2.34000 | 6 | 14 |
| B |  |  |  |  |
| B | C | 2.28167 | 6 | 23 |
|  | C |  |  |  |
| D | C | 2.25833 | 6 | 24 |
| D |  |  |  |  |
| D | C E | 2.23500 | 6 | 20 |
| D | C E |  |  |  |
| D | C E | 2.23167 | 6 | 17 |
| D | C E |  |  |  |
| D | C E | 2.22333 | 6 | 26 |
| D | C E |  |  |  |
| D | C E | 2.21500 | 6 | 30 |
| D | E |  |  |  |
| D | F E | 2.20167 | 6 | 27 |
| D | F E |  |  |  |
| D | F E | 2.19833 | 6 | 22 |
| D | F E |  |  |  |
| D | G F E | 2.19333 | 6 | 16 |
|  | G F E |  |  |  |
|  | G F E | 2.18500 | 6 | 31 |
|  | G F E |  |  |  |
|  | G F E | 2.17000 | 6 | 7 |
|  | G F |  |  |  |
| H | G F | 2.13167 | 6 | 12 |
| H | G |  |  |  |
| H | G I | 2.12167 | 6 | 28 |
| H | I |  |  |  |
| H | I J | 2.07667 | 6 | 8 |
| H | I J |  |  |  |
| H | I J | 2.06000 | 6 | 10 |
|  | I J |  |  |  |
|  | I J | 2.05833 | 6 | 19 |
|  | I J |  |  |  |
|  | I J | 2.05500 | 6 | 29 |
|  | J |  |  |  |

The GLM Procedure

t Tests (LSD) for SD\_cm\_

Means with the same letter are not significantly different.

| t Grouping |  | Mean | N | Genotype |
| --- | --- | --- | --- | --- |
|  | K J | 2.04333 | 6 | 13 |
|  | K J |  |  |  |
| L | K J | 2.02833 | 6 | 21 |
| L | K |  |  |  |
| L | K M | 1.98167 | 6 | 11 |
| L | M |  |  |  |
| L | M | 1.96000 | 6 | 1 |
|  | M |  |  |  |
|  | M | 1.95333 | 6 | 5 |
|  | M |  |  |  |
|  | M | 1.95167 | 6 | 6 |
|  | N | 1.87167 | 6 | 9 |
|  | N |  |  |  |
| O | N | 1.82500 | 6 | 3 |
| O | N |  |  |  |
| O | N | 1.81500 | 6 | 4 |
| O |  |  |  |  |
| O | P | 1.79167 | 6 | 25 |
|  | P |  |  |  |
|  | P | 1.72667 | 6 | 2 |
|  | Q | 1.02333 | 6 | 18 |

14:00 Thursday, February 18, 2026 24

The GLM Procedure

t Tests (LSD) for LA\_\_cm2\_

NOTE: This test controls the Type I comparisonwise error rate, not the experimentwise error rate.

|  |  |
| --- | --- |
| Alpha | 0.05 |
| Error Degrees of Freedom | 124 |
| Error Mean Square | 268.3265 |
| Critical Value of t | 1.97928 |
| Least Significant Difference | 18.719 |

Means with the same letter are not significantly different.

| t Grouping |  | Mean | N | Genotype |
| --- | --- | --- | --- | --- |
|  | A | 436.687 | 6 | 23 |
|  | A |  |  |  |
|  | A | 432.578 | 6 | 17 |
|  | B | 412.817 | 6 | 22 |
|  | B |  |  |  |
|  | B | 408.780 | 6 | 21 |
|  | B |  |  |  |
|  | B | 408.272 | 6 | 30 |
|  | B |  |  |  |
|  | B | 405.885 | 6 | 24 |
|  | C | 378.510 | 6 | 14 |
|  | C |  |  |  |
| D | C | 369.615 | 6 | 5 |
| D |  |  |  |  |
| D | C | 365.462 | 6 | 8 |
| D |  |  |  |  |
| D | C | 361.985 | 6 | 26 |
| D |  |  |  |  |
| D | E | 354.625 | 6 | 6 |
| D |  |  |  |  |
| D | E | 354.045 | 6 | 7 |
| D |  |  |  |  |
| D | E | 353.438 | 6 | 15 |
|  | E |  |  |  |
| F | E | 341.818 | 6 | 20 |
| F |  |  |  |  |
| F | G | 333.835 | 6 | 25 |
| F |  |  |  |  |
| F | G | 328.942 | 6 | 11 |
| F |  |  |  |  |
| F | G | 324.165 | 6 | 16 |
|  | I |  |  |  |
| J | I | 317.643 | 6 | 4 |
| J |  |  |  |  |
| J | I | 314.092 | 6 | 9 |
| J |  |  |  |  |
| J | I |  |  |  |

14:00 Thursday, February 18, 2026 25

The GLM Procedure

t Tests (LSD) for LA\_\_cm2\_

Means with the same letter are not significantly different.

| t Grouping |  | Mean | N | Genotype |
| --- | --- | --- | --- | --- |

|  |  |  |  |  |  |  |
| --- | --- | --- | --- | --- | --- | --- |
| J | I | K | H | 310.385 | 6 | 31 |
| J | I | K | H |  |  |  |
| J | I | K | H | 310.367 | 6 | 27 |
| J | I | K |  |  |  |  |
| J | I | K |  | 309.133 | 6 | 28 |
| J | I | K |  |  |  |  |
| J | I | K |  | 307.922 | 6 | 10 |
| J |  | K |  |  |  |  |
| J |  | K | L | 303.715 | 6 | 13 |
|  |  | K | L |  |  |  |
| M |  | K | L | 293.902 | 6 | 12 |
| M |  |  | L |  |  |  |
| M |  | N | L | 287.732 | 6 | 2 |
| M |  | N | L |  |  |  |
| M |  | N | L | 286.902 | 6 | 29 |
| M |  | N |  |  |  |  |
| M |  | N |  | 284.977 | 6 | 1 |
|  |  | N |  |  |  |  |
|  |  | N |  | 274.765 | 6 | 3 |
|  |  | N |  |  |  |  |
|  |  | N |  | 269.915 | 6 | 19 |
|  |  | O |  | 104.772 | 6 | 18 |

14:00 Thursday, February 18, 2026 26

The GLM Procedure

t Tests (LSD) for PFW\_\_g\_\_

NOTE: This test controls the Type I comparisonwise error rate, not the experimentwise error rate.

|  |  |
| --- | --- |
| Alpha | 0.05 |
| Error Degrees of Freedom | 124 |
| Error Mean Square | 0.460161 |
| Critical Value of t | 1.97928 |
| Least Significant Difference | 0.7752 |

Means with the same letter are not significantly different.

| t Grouping |  | Mean | N | Genotype |
| --- | --- | --- | --- | --- |
|  | A | 21.5000 | 6 | 23 |
|  | A |  |  |  |
| B | A | 20.8167 | 6 | 22 |
| B |  |  |  |  |
| B |  | 20.5333 | 6 | 14 |
| B |  |  |  |  |
| B | C | 20.1667 | 6 | 17 |
| B | C |  |  |  |
| B | C | 20.0500 | 6 | 24 |
|  | C |  |  |  |
|  | C | 19.4167 | 6 | 27 |
|  | D | 18.5167 | 6 | 30 |
|  | D |  |  |  |
|  | D | 18.4333 | 6 | 26 |
|  | D |  |  |  |
| E | D | 18.2167 | 6 | 5 |
| E | D |  |  |  |
| E | D | 18.0667 | 6 | 31 |
| E | D |  |  |  |
| E | D | 18.0167 | 6 | 28 |
| E | D | F |  |  |
| E | D | F |  |  |
| E | G | 17.8333 | 6 | 15 |
| E | G |  |  |  |
| E | G | 17.5500 | 6 | 16 |
|  | G |  |  |  |
|  | G | 17.2833 | 6 | 7 |
|  | G |  |  |  |
|  | G | 17.2167 | 6 | 20 |
|  | H |  |  |  |
| I | H | 16.6500 | 6 | 21 |
| I |  |  |  |  |
| I | J | 16.2667 | 6 | 8 |
| I | J |  |  |  |
| I | J | 16.0500 | 6 | 19 |
|  | J | K |  |  |
| L | J | 15.6000 | 6 | 6 |
|  | K |  |  |  |

L

K

14:00 Thursday, February 18, 2026 27

#### The GLM Procedure

t Tests (LSD) for PFW\_g\_

Means with the same letter are not significantly different.

| t Grouping |  |  | Mean | N | Genotype |
| --- | --- | --- | --- | --- | --- |
| L | M | K | 15.3500 | 6 | 4 |
| L | M |  |  |  |  |
| L | M |  | 15.2167 | 6 | 13 |
| L | M |  |  |  |  |
| L | M |  | 14.8333 | 6 | 12 |
|  | M |  |  |  |  |
| N | M |  | 14.5833 | 6 | 1 |
| N |  |  |  |  |  |
| N | O |  | 13.9833 | 6 | 3 |
|  | O |  |  |  |  |
| P | O |  | 13.7500 | 6 | 9 |
| P | O |  |  |  |  |
| P | O |  | 13.6333 | 6 | 10 |
| P | O |  |  |  |  |
| P | O | Q | 13.4000 | 6 | 2 |
| P |  | Q |  |  |  |
| P |  | Q | 13.1333 | 6 | 25 |
| P |  | Q |  |  |  |
| P |  | Q | 13.0167 | 6 | 11 |
|  |  | Q |  |  |  |
|  |  | Q | 12.8500 | 6 | 29 |
|  | R |  | 4.3000 | 6 | 18 |

14:00 Thursday, February 18, 2026 28

#### The GLM Procedure

t Tests (LSD) for RFW\_g\_

NOTE: This test controls the Type I comparisonwise error rate, not the experimentwise error rate.

|  |  |
| --- | --- |
| Alpha | 0.05 |
| Error Degrees of Freedom | 124 |
| Error Mean Square | 0.398503 |
| Critical Value of t | 1.97928 |
| Least Significant Difference | 0.7214 |

Means with the same letter are not significantly different.

| t Grouping |  |  | Mean | N | Genotype |
| --- | --- | --- | --- | --- | --- |
|  | A |  | 10.7167 | 6 | 30 |
|  | B |  | 8.0500 | 6 | 19 |
|  | B |  |  |  |  |
|  | B |  | 7.8500 | 6 | 27 |
|  | B |  |  |  |  |
| C | B |  | 7.5833 | 6 | 28 |
| C |  |  |  |  |  |
| C | D |  | 7.0083 | 6 | 4 |
|  | D |  |  |  |  |
| E | D |  | 6.8500 | 6 | 17 |
| E | D |  |  |  |  |
| E | D |  | 6.8383 | 6 | 13 |
| E | D |  |  |  |  |
| E | D |  | 6.7800 | 6 | 1 |
| E | D |  |  |  |  |
| E | D |  | 6.5667 | 6 | 23 |
| E | D |  |  |  |  |
| E | D |  | 6.4167 | 6 | 20 |
| E | D |  |  |  |  |
| E | D |  | 6.3833 | 6 | 21 |
| E |  |  |  |  |  |
| E |  |  | 6.2550 | 6 | 3 |
| E |  |  |  |  |  |
| E | F |  | 6.2000 | 6 | 14 |
|  | F |  |  |  |  |

|  |  |  |  |  |
| --- | --- | --- | --- | --- |
| G | F | 5.5167 | 6 | 24 |
| G |  |  |  |  |
| G | H | 5.3333 | 6 | 22 |
| G | H |  |  |  |
| G | H | 5.3167 | 6 | 16 |
| G | H |  |  |  |
| G | H | 5.2800 | 6 | 6 |
| G | H |  |  |  |
| G | H | 5.2667 | 6 | 15 |
| G | H |  |  |  |
| G | H | 5.2583 | 6 | 9 |
| G | H |  |  |  |

14:00 Thursday, February 18, 2026 29

### The GLM Procedure

#### t Tests (LSD) for RFW\_\_g\_\_

Means with the same letter are not significantly different.

| t Grouping |  | Mean | N | Genotype |
| --- | --- | --- | --- | --- |
| G | H | 5.1167 | 6 | 29 |
| G | H |  |  |  |
| G | H | 5.1000 | 6 | 31 |
| G | H |  |  |  |
| G | H | 5.0450 | 6 | 5 |
| G | H |  |  |  |
| G | H | 5.0167 | 6 | 2 |
| G | H |  |  |  |
| G | H I | 4.9333 | 6 | 26 |
| G | H I |  |  |  |
| J | H I | 4.8583 | 6 | 8 |
| J | H I |  |  |  |
| J | H I | 4.6717 | 6 | 11 |
| J | H I |  |  |  |
| J | H I | 4.6550 | 6 | 7 |
| J | I |  |  |  |
| J | K I | 4.2867 | 6 | 12 |
| J | K |  |  |  |
| J | K | 4.2000 | 6 | 25 |
|  | K |  |  |  |
|  | K | 3.8467 | 6 | 10 |
|  | L | 1.5000 | 6 | 18 |

14:00 Thursday, February 18, 2026 30

### The GLM Procedure

#### t Tests (LSD) for PDW\_\_g\_\_

NOTE: This test controls the Type I comparisonwise error rate, not the experimentwise error rate.

|  |  |
| --- | --- |
| Alpha | 0.05 |
| Error Degrees of Freedom | 124 |
| Error Mean Square | 0.009234 |
| Critical Value of t | 1.97928 |
| Least Significant Difference | 0.1098 |

Means with the same letter are not significantly different.

| t Grouping |  | Mean | N | Genotype |
| --- | --- | --- | --- | --- |
|  | A | 2.00500 | 6 | 21 |
|  | A |  |  |  |
| B | A | 1.96333 | 6 | 20 |
| B |  |  |  |  |
| B | C | 1.88667 | 6 | 23 |
| B | C |  |  |  |
| B | C | 1.87500 | 6 | 22 |
| B | C |  |  |  |
| B | C | 1.85500 | 6 | 30 |
|  | C |  |  |  |
|  | C | 1.84833 | 6 | 27 |
|  | C |  |  |  |
| D | C | 1.79333 | 6 | 17 |
| D |  |  |  |  |
| D | E | 1.69333 | 6 | 14 |

|  |  |  |  |  |  |
| --- | --- | --- | --- | --- | --- |
| F |  | E |  |  |  |
| F |  | E |  | 1.67833 | 6 24 |
| F |  | E |  |  |  |
| F |  | E |  | 1.62500 | 6 26 |
| F |  |  |  |  |  |
| F |  | G |  | 1.58000 | 6 28 |
|  |  | G |  |  |  |
| H |  | G |  | 1.49667 | 6 16 |
| H |  |  |  |  |  |
| H |  | I |  | 1.47000 | 6 4 |
| H |  | I |  |  |  |
| H |  | I |  | 1.46167 | 6 5 |
| H |  | I |  |  |  |
| H |  | I | J | 1.44667 | 6 15 |
| H |  | I | J |  |  |
| H | K | I | J | 1.41667 | 6 31 |
|  | K | I | J |  |  |
|  | K | I | J | 1.37000 | 6 7 |
|  | K |  | J |  |  |
|  | K | L | J | 1.34000 | 6 19 |
|  | K | L |  |  |  |
|  | K | L |  | 1.32833 | 6 8 |
|  | K | L |  |  |  |

14:00 Thursday, February 18, 2026 31

###### The GLM Procedure

###### t Tests (LSD) for PDW\_\_g\_\_

Means with the same letter are not significantly different.

| t Grouping |  |  | Mean | N | Genotype |
| --- | --- | --- | --- | --- | --- |
| N | K | L | 1.31167 | 6 | 29 |
|  |  | L |  |  |  |
|  |  | L | 1.24833 | 6 | 1 |
|  |  | L |  |  |  |
|  |  | L | 1.24000 | 6 | 3 |
|  | M |  | 1.10667 | 6 | 2 |
|  | M |  | 1.04333 | 6 | 6 |
|  | M |  |  |  |  |
|  | M | 0 | 1.00500 | 6 | 9 |
| N |  | 0 |  |  |  |
| N |  | 0 | 0.98833 | 6 | 25 |
| N |  | 0 |  |  |  |
| N |  | 0 | 0.92333 | 6 | 12 |
|  |  | 0 |  |  |  |
|  |  | 0 | 0.91667 | 6 | 13 |
|  | P |  | 0.79167 | 6 | 11 |
|  | Q |  | 0.67667 | 6 | 10 |
|  | R |  | 0.44833 | 6 | 18 |

14:00 Thursday, February 18, 2026 32

###### The GLM Procedure

###### t Tests (LSD) for RDW\_\_g\_\_

NOTE: This test controls the Type I comparisonwise error rate, not the experimentwise error rate.

|  |  |
| --- | --- |
| Alpha | 0.05 |
| Error Degrees of Freedom | 124 |
| Error Mean Square | 0.00157 |
| Critical Value of t | 1.97928 |
| Least Significant Difference | 0.0453 |

Means with the same letter are not significantly different.

| t Grouping |  |  | Mean | N | Genotype |
| --- | --- | --- | --- | --- | --- |
|  | A |  | 0.63667 | 6 | 26 |
|  | A |  |  |  |  |
|  | A |  | 0.62333 | 6 | 30 |
|  | A |  |  |  |  |

|  |  |  |  |  |  |  |  |
| --- | --- | --- | --- | --- | --- | --- | --- |
|  | B |  | A |  | 0.59500 | 6 | 27 |
|  | B |  |  |  |  |  |  |
|  | B |  |  |  | 0.57500 | 6 | 17 |
|  |  |  | C |  | 0.51500 | 6 | 19 |
|  |  |  | D |  | 0.45000 | 6 | 28 |
|  |  |  | D |  |  |  |  |
|  |  |  | D |  | 0.44333 | 6 | 21 |
|  |  |  | D |  |  |  |  |
|  | E |  | D |  | 0.43500 | 6 | 20 |
|  | E |  | D |  |  |  |  |
|  | E |  | D |  | 0.42833 | 6 | 29 |
|  | E |  |  |  |  |  |  |
|  | E |  |  |  | 0.39333 | 6 | 5 |
|  | E |  |  |  |  |  |  |
|  | E |  |  |  | 0.39000 | 6 | 4 |
|  |  |  | F |  | 0.34333 | 6 | 15 |
|  |  |  | F |  |  |  |  |
|  |  |  | F |  | 0.34167 | 6 | 16 |
|  |  |  | F |  |  |  |  |
|  | G |  | F |  | 0.33500 | 6 | 22 |
|  | G |  | F |  |  |  |  |
|  | G |  | F |  | 0.33167 | 6 | 23 |
|  | G |  | F |  |  |  |  |
|  | G |  | F |  | 0.32667 | 6 | 13 |
|  | G |  | F |  |  |  |  |
|  | G |  | F | H | 0.32167 | 6 | 14 |
|  | G |  | F | H |  |  |  |
|  | G | I | F | H | 0.31667 | 6 | 1 |
|  | G | I | F | H |  |  |  |
| J | G | I | F | H | 0.30333 | 6 | 6 |
| J | G | I |  | H |  |  |  |

14:00 Thursday, February 18, 2026 33

14:00 Thursday, February 18, 2026 33

###### The GLM Procedure

###### t Tests (LSD) for RDW\_\_g\_\_

Means with the same letter are not significantly different.

| t Grouping |  |  |  |  | Mean | N | Genotype |
| --- | --- | --- | --- | --- | --- | --- | --- |
| J | G | I | K | H | 0.29333 | 6 | 3 |
| J |  | I | K | H |  |  |  |
| J |  | I | K | H | 0.27833 | 6 | 2 |
| J |  | I | K |  |  |  |  |
| J |  | I | K |  | 0.27333 | 6 | 31 |
| J |  | I | K |  |  |  |  |
| J |  | I | K |  | 0.27167 | 6 | 8 |
| J |  |  | K |  |  |  |  |
| J |  |  | K |  | 0.26833 | 6 | 7 |
| J |  |  | K |  |  |  |  |
| J | L |  | K |  | 0.25833 | 6 | 24 |
|  | L |  | K |  |  |  |  |
|  | L |  | K |  | 0.25000 | 6 | 11 |
|  | L |  | K |  |  |  |  |
|  | L |  | K | M | 0.24833 | 6 | 12 |
|  | L |  |  | M |  |  |  |
|  | L |  | N | M | 0.22167 | 6 | 9 |
|  |  |  | N | M |  |  |  |
|  |  |  | N | M | 0.20333 | 6 | 25 |
|  |  |  | N |  |  |  |  |
|  |  |  | N |  | 0.19167 | 6 | 10 |
|  |  |  | O |  | 0.10500 | 6 | 18 |

14:00 Thursday, February 18, 2026 34

###### The GLM Procedure

###### t Tests (LSD) for EC\_\_\_\_\_

NOTE: This test controls the Type I comparisonwise error rate, not the experimentwise error rate.

|  |  |
| --- | --- |
| Alpha | 0.05 |
| Error Degrees of Freedom | 124 |
| Error Mean Square | 12.79635 |
| Critical Value of t | 1.97928 |
| Least Significant Difference | 4.0878 |

| t Grouping |  |  | Mean | N | Genotype |  |
| --- | --- | --- | --- | --- | --- | --- |
|  |  | A | 49.181 | 6 | 20 |  |
|  |  | B | 43.341 | 6 | 10 |  |
|  |  | B |  |  |  |  |
|  |  | B | 42.588 | 6 | 15 |  |
|  |  | B |  |  |  |  |
|  |  | B | 42.142 | 6 | 9 |  |
|  |  | B |  |  |  |  |
|  |  | B | 41.360 | 6 | 18 |  |
|  |  | B |  |  |  |  |
|  |  | B | 41.034 | 6 | 7 |  |
|  |  | B |  |  |  |  |
| C |  | B | 40.255 | 6 | 23 |  |
| C |  | B |  |  |  |  |
| C |  | B | 40.208 | 6 | 22 |  |
| C |  | B |  |  |  |  |
| C |  | B | D | 40.013 | 6 | 30 |
| C |  |  | D |  |  |  |
|  |  | E | D | 36.808 | 6 | 12 |
|  |  | E | D |  |  |  |
| F |  | E | D | 36.048 | 6 | 16 |
| F |  | E |  |  |  |  |
| F |  | E |  | 35.806 | 6 | 14 |
| F |  |  |  |  |  |  |
|  |  | G | 32.485 | 6 | 2 |  |
|  |  | G |  |  |  |  |
| H |  | G | 30.693 | 6 | 25 |  |
| H |  | G |  |  |  |  |
| H |  | G | I | 30.529 | 6 | 19 |
| H |  | G | I |  |  |  |
| H |  | G | I | 30.434 | 6 | 11 |
| H |  | G | I |  |  |  |
| H | J | G | I | 29.096 | 6 | 28 |
| H | J | G | I |  |  |  |
| H | J | G | I | 28.434 | 6 | 21 |
| H | J |  | I |  |  |  |
| H | J |  | I | 28.197 | 6 | 13 |
| H | J |  | I |  |  |  |

14:00 Thursday, February 18, 2026 35

14:00 Thursday, February 18, 2026 35

#### The GLM Procedure

t Tests (LSD) for EC\_\_\_\_\_

Means with the same letter are not significantly different.

| t Grouping |  |  |  | Mean | N | Genotype |
| --- | --- | --- | --- | --- | --- | --- |
| H | J | K | I | 27.456 | 6 | 4 |
| H | J | K | I |  |  |  |
| H | J | K | I | 27.385 | 6 | 29 |
| H | J | K | I |  |  |  |
| H | J | K | I | 27.293 | 6 | 17 |
|  | J | K | I |  |  |  |
| L | J | K | I | 26.601 | 6 | 31 |
| L | J | K |  |  |  |  |
| L | J | K | M | 25.160 | 6 | 26 |
| L |  | K | M |  |  |  |
| L | N | K | M | 24.044 | 6 | 5 |
| L | N | K | M |  |  |  |
| L | N | K | M | 23.776 | 6 | 24 |
| L | N |  | M |  |  |  |
| L | N | O | M | 22.631 | 6 | 3 |
|  | N | O | M |  |  |  |
|  | N | O | M | 22.369 | 6 | 8 |
|  | N | O |  |  |  |  |
|  | N | O |  | 20.178 | 6 | 27 |
|  | N | O |  |  |  |  |
|  | N | O |  | 20.133 | 6 | 6 |
|  |  | O |  |  |  |  |
|  |  | O |  | 19.606 | 6 | 1 |

14:00 Thursday, February 18, 2026 36

#### The GLM Procedure

### t Tests (LSD) for LRWC\_\_\_\_

NOTE: This test controls the Type I comparisonwise error rate, not the experimentwise error rate.

|  |  |
| --- | --- |
| Alpha | 0.05 |
| Error Degrees of Freedom | 124 |
| Error Mean Square | 33.11233 |
| Critical Value of t | 1.97928 |
| Least Significant Difference | 6.5757 |

Means with the same letter are not significantly different.

| t Grouping |  |  |  |  | Mean | N | Genotype |  |
| --- | --- | --- | --- | --- | --- | --- | --- | --- |
| B |  | A |  |  | 75.560 | 6 | 19 |  |
|  |  | A |  |  |  |  |  |  |
|  |  | A |  |  | 73.674 | 6 | 7 |  |
|  |  | A |  |  |  |  |  |  |
|  |  | A | C |  | 70.546 | 6 | 4 |  |
|  |  | A | C |  |  |  |  |  |
|  |  | A | C |  | 70.502 | 6 | 28 |  |
|  |  | A | C |  |  |  |  |  |
|  |  | A | C |  | 70.317 | 6 | 2 |  |
|  |  | A | C |  |  |  |  |  |
|  |  | A | C |  | 70.024 | 6 | 13 |  |
|  |  | A | C |  |  |  |  |  |
|  |  | D | A | C | 69.393 | 6 | 21 |  |
|  |  | D |  | C |  |  |  |  |
|  |  | D | E | C | 68.472 | 6 | 9 |  |
|  |  | D | E | C |  |  |  |  |
|  |  | D | E | C | 68.282 | 6 | 29 |  |
|  |  | D | E | C |  |  |  |  |
|  |  | D | E | C | 68.124 | 6 | 23 |  |
|  |  | D | E | C |  |  |  |  |
| D | E | C | 68.032 | 6 | 3 |  |  |  |
| D | E | C |  |  |  |  |  |  |
| F | B | D | E | C | 67.985 | 6 | 31 |  |
| F | B | D | E | C |  |  |  |  |
| F | B | D | E | C | G | 67.901 | 6 | 26 |
| F | B | D | E | C | G |  |  |  |
| F | B | D | E | C | G | 67.513 | 6 | 14 |
| F |  | D | E | C | G |  |  |  |
| F |  | D | E | C | G | 66.666 | 6 | 10 |
| F |  | D | E | C | G |  |  |  |
| F |  | D | E | C | G | 66.434 | 6 | 6 |
| F |  | D | E | C | G |  |  |  |
| F |  | D | E | C | G | 66.406 | 6 | 25 |
| F |  | D | E | C | G |  |  |  |
| F |  | D | E | C | G | 66.222 | 6 | 15 |
| F |  | D | E | C | G |  |  |  |
| F |  | D | E | C | G | 65.947 | 6 | 30 |
| F |  | D | E | C | G |  |  |  |

14:00 Thursday, February 18, 2026 37

#### The GLM Procedure

##### t Tests (LSD) for LRWC\_\_\_\_

Means with the same letter are not significantly different.

| t Grouping |  |  |  |  | Mean | N | Genotype |  |
| --- | --- | --- | --- | --- | --- | --- | --- | --- |
| F |  | D | E | C | G | 65.887 | 6 | 12 |
| F |  | D | E | C | G |  |  |  |
| F |  | D | E | C | G | 65.606 | 6 | 1 |
| F |  | D | E | C | G |  |  |  |
| F |  | D | E | C | G | 65.203 | 6 | 27 |
| F |  | D | E | C | G |  |  |  |
| F |  | D | E | C | G | 65.089 | 6 | 18 |
| F |  | D | E | C | G |  |  |  |
| F |  | D | E | C | G | 64.562 | 6 | 16 |
| F |  | D | E | C | G |  |  |  |
| F |  | D | E | C | G | 64.362 | 6 | 5 |
| F |  | D | E |  | G |  |  |  |
| F | H | D | E |  | G | 63.191 | 6 | 11 |
| F | H |  | E |  | G |  |  |  |
| F | H |  | E |  | G | 62.745 | 6 | 22 |

|  |  |  |  |  |  |  |
| --- | --- | --- | --- | --- | --- | --- |
| F | H | E | G | 62.524 | 6 | 17 |
| F | H |  | G |  |  |  |
| F | H |  | G | 61.441 | 6 | 24 |
| F | H |  | G |  |  |  |
|  | H |  | G | 61.378 | 6 | 8 |
|  | H |  |  |  |  |  |
|  | H |  |  | 56.793 | 6 | 20 |

14:00 Thursday, February 18, 2026 38

The GLM Procedure

t Tests (LSD) for Chla\_\_mg\_g\_1\_

NOTE: This test controls the Type I comparisonwise error rate, not the experimentwise error rate.

|  |  |
| --- | --- |
| Alpha | 0.05 |
| Error Degrees of Freedom | 124 |
| Error Mean Square | 0.120174 |
| Critical Value of t | 1.97928 |
| Least Significant Difference | 0.3961 |

Means with the same letter are not significantly different.

| t Grouping |  |  |  | Mean | N | Genotype |
| --- | --- | --- | --- | --- | --- | --- |
|  |  | A |  | 5.7133 | 6 | 11 |
|  |  | A |  |  |  |  |
| B |  | A |  | 5.5489 | 6 | 10 |
| B |  | A |  |  |  |  |
| B |  | A | C | 5.4780 | 6 | 16 |
| B |  |  | C |  |  |  |
| B |  | D | C | 5.2608 | 6 | 12 |
| B |  | D | C |  |  |  |
| B | E | D | C | 5.2121 | 6 | 19 |
|  | E | D | C |  |  |  |
| F | E | D | C | 5.1473 | 6 | 13 |
| F | E | D | C |  |  |  |
| F | E | D | C | 5.1154 | 6 | 31 |
| F | E | D | C |  |  |  |
| F | E | D | C | 5.1138 | 6 | 9 |
| F | E | D |  |  |  |  |
| F | E | D |  | 5.0175 | 6 | 15 |
| F | E | D |  |  |  |  |
| F | E | D |  | 4.9949 | 6 | 18 |
| F | E |  |  |  |  |  |
| F | E | G |  | 4.8298 | 6 | 14 |
| F |  | G |  |  |  |  |
| F |  | G |  | 4.7972 | 6 | 1 |
| F |  | G |  |  |  |  |
| F |  | G | H | 4.7525 | 6 | 3 |
|  |  | G | H |  |  |  |
| I |  | G | H | 4.5470 | 6 | 8 |
| I |  | G | H |  |  |  |
| I | J | G | H | 4.4654 | 6 | 29 |
| I | J |  | H |  |  |  |
| I | J | K | H | 4.3633 | 6 | 20 |
| I | J | K |  |  |  |  |
| I | J | K |  | 4.2617 | 6 | 30 |
| I | J | K |  |  |  |  |
| I | J | K |  | 4.2572 | 6 | 7 |
|  | J | K |  |  |  |  |
| L | J | K |  | 4.1446 | 6 | 17 |
| L | J | K |  |  |  |  |

14:00 Thursday, February 18, 2026 39

The GLM Procedure

t Tests (LSD) for Chla\_\_mg\_g\_1\_

Means with the same letter are not significantly different.

| t Grouping |  |  |  | Mean | N | Genotype |
| --- | --- | --- | --- | --- | --- | --- |
| L | J | K |  | 4.1038 | 6 | 22 |
| L |  | K |  |  |  |  |
| L |  | K | M | 4.0649 | 6 | 6 |
| L |  | K | M |  |  |  |
| L |  | K | M | 4.0607 | 6 | 23 |
| L |  | K | M |  |  |  |
| L | N | K | M | 4.0155 | 6 | 4 |
| L | N | K | M |  |  |  |
| L | N | K | M | 4.0006 | 6 | 21 |
| L | N |  | M |  |  |  |
| L | N | O | M | 3.8600 | 6 | 27 |
| L | N | O | M |  |  |  |
| L | N | O | M | 3.7903 | 6 | 24 |
| L | N | O | M |  |  |  |
| L | N | O | M | 3.7518 | 6 | 28 |
|  | N | O | M |  |  |  |
|  | N | O | M | 3.6759 | 6 | 25 |
|  | N | O |  |  |  |  |
|  | N | O |  | 3.6315 | 6 | 26 |
|  |  | O |  |  |  |  |
|  |  | O |  | 3.5313 | 6 | 5 |
|  |  | O |  |  |  |  |
|  |  | O |  | 3.5019 | 6 | 2 |

14:00 Thursday, February 18, 2026 40

The GLM Procedure

t Tests (LSD) for Chlb\_\_mg\_g\_1\_

NOTE: This test controls the Type I comparisonwise error rate, not the experimentwise error rate.

|  |  |
| --- | --- |
| Alpha | 0.05 |
| Error Degrees of Freedom | 124 |
| Error Mean Square | 0.025918 |
| Critical Value of t | 1.97928 |
| Least Significant Difference | 0.184 |

Means with the same letter are not significantly different.

| t Grouping |  | Mean | N | Genotype |
| --- | --- | --- | --- | --- |
|  | A | 3.99776 | 6 | 18 |
|  | B | 3.75613 | 6 | 1 |
|  | C | 3.44400 | 6 | 10 |
|  | C |  |  |  |
| D | C | 3.39300 | 6 | 4 |
| D |  |  |  |  |
| D | E | 3.21988 | 6 | 22 |
|  | E |  |  |  |
| F | E | 3.11833 | 6 | 31 |
| F | E |  |  |  |
| F | E | 3.08986 | 6 | 11 |
| F | G | 2.99550 | 6 | 3 |
|  | G |  |  |  |
| H | G | 2.90347 | 6 | 30 |
| H | G |  |  |  |
| H | G | 2.89811 | 6 | 19 |
| H | G |  |  |  |
| H | G | 2.86446 | 6 | 17 |
| H | G |  |  |  |
| H | G | I | 6 | 13 |
| H | G | I |  |  |
| H | G | I | 6 | 8 |
| H |  | I |  |  |
| H |  | I | 6 | 26 |
| H |  | I |  |  |
| H |  | I | 6 | 7 |
| H |  | I |  |  |
| H |  | I | 6 | 23 |
|  |  | I |  |  |
|  |  | I | 6 | 20 |
|  | J | 2.29025 | 6 | 9 |
|  | J |  |  |  |
| K | J | 2.27631 | 6 | 28 |
| K | J |  |  |  |

14:00 Thursday, February 18, 2026 41

The GLM Procedure

t Tests (LSD) for Chlb\_\_mg\_g\_1\_

Means with the same letter are not significantly different.

| t Grouping |  | Mean | N | Genotype |
| --- | --- | --- | --- | --- |
| K | J | 2.21762 | 6 | 29 |
| K | J |  |  |  |
| K | J | 2.20921 | 6 | 21 |
| K | J |  |  |  |
| K | J | 2.19026 | 6 | 15 |
| K | J |  |  |  |
| K | J | 2.18358 | 6 | 27 |
| K | J |  |  |  |
| K | J | 2.15552 | 6 | 24 |
| K | J |  |  |  |
| K | J | 2.12711 | 6 | 5 |
| K | J |  |  |  |
| K | J | 2.11588 | 6 | 14 |
| K |  |  |  |  |
| K |  | 2.09756 | 6 | 6 |
|  |  | 2.01450 | 6 | 2 |
|  | M |  |  |  |
|  | M |  |  |  |
|  | M | 1.90558 | 6 | 25 |
|  | N | 1.71107 | 6 | 12 |
|  | O | 1.50992 | 6 | 16 |

14:00 Thursday, February 18, 2026 42

The GLM Procedure

t Tests (LSD) for Total\_Ch1\_\_mg\_g\_1\_

NOTE: This test controls the Type I comparisonwise error rate, not the experimentwise error rate.

|  |  |
| --- | --- |
| Alpha | 0.05 |
| Error Degrees of Freedom | 124 |
| Error Mean Square | 0.170501 |
| Critical Value of t | 1.97928 |
| Least Significant Difference | 0.4719 |

Means with the same letter are not significantly different.

| t Grouping |  | Mean | N | Genotype |
| --- | --- | --- | --- | --- |
|  | A | 8.9923 | 6 | 10 |
|  | A |  |  |  |
|  | A | 8.9917 | 6 | 18 |
|  | A |  |  |  |
|  | A | 8.8024 | 6 | 11 |
|  | A |  |  |  |
| B | A | 8.5528 | 6 | 1 |
| B |  |  |  |  |
| B | C | 8.2328 | 6 | 31 |
| B | C |  |  |  |
| B | C | 8.1089 | 6 | 19 |
|  | C |  |  |  |
|  | C | 7.9872 | 6 | 13 |
|  | D |  |  |  |
|  | E | 7.7466 | 6 | 3 |
|  | E |  |  |  |
| F | E | 7.4077 | 6 | 4 |
| F | E |  |  |  |
| F | E | 7.4034 | 6 | 9 |
| F | E |  |  |  |
| F | E | 7.3796 | 6 | 8 |
| F | E |  |  |  |
| F | E | 7.3233 | 6 | 22 |
| F |  |  |  |  |
| F | G | 7.2069 | 6 | 15 |
| F | G |  |  |  |
| F | G | 7.1645 | 6 | 30 |
| F | G |  |  |  |
| F | G | 7.0446 | 6 | 7 |
| F | G |  |  |  |
| F | G | 7.0253 | 6 | 20 |

|  |  |  |  |  |  |
| --- | --- | --- | --- | --- | --- |
| F | G | H |  |  |  |
| F | G | H | 7.0077 | 6 | 17 |
| F | G | H |  |  |  |
| F | G | H | 6.9763 | 6 | 16 |
| F | G | H |  |  |  |
| F | G | H | 6.9713 | 6 | 12 |
| F | G | H |  |  |  |

14:00 Thursday, February 18, 2026 43

###### The GLM Procedure

t Tests (LSD) for Total\_Ch1\_\_mg\_g\_1\_

Means with the same letter are not significantly different.

| t Grouping |  |  | Mean | N | Genotype |
| --- | --- | --- | --- | --- | --- |
| F | G | H | 6.9454 | 6 | 14 |
|  | G | H |  |  |  |
| I | G | H | 6.8481 | 6 | 23 |
| I |  | H |  |  |  |
| I |  | H | 6.6821 | 6 | 29 |
| I |  |  |  |  |  |
| I | J |  | 6.4290 | 6 | 26 |
|  | J |  |  |  |  |
| K | J |  | 6.2091 | 6 | 21 |
| K | J |  |  |  |  |
| K | J |  | 6.1618 | 6 | 6 |
| K | J |  |  |  |  |
| K | J | L | 6.0425 | 6 | 27 |
| K | J | L |  |  |  |
| K | J | L | 6.0277 | 6 | 28 |
| K |  | L |  |  |  |
| K | M | L | 5.9451 | 6 | 24 |
|  | M | L |  |  |  |
|  | M | L | 5.6577 | 6 | 5 |
|  | M | L |  |  |  |
|  | M | L | 5.5808 | 6 | 25 |
|  | M |  |  |  |  |
|  | M |  | 5.5156 | 6 | 2 |

14:00 Thursday, February 18, 2026 44

###### The GLM Procedure

t Tests (LSD) for Car\_\_mg\_g\_1\_

NOTE: This test controls the Type I comparisonwise error rate, not the experimentwise error rate.

|  |  |
| --- | --- |
| Alpha | 0.05 |
| Error Degrees of Freedom | 124 |
| Error Mean Square | 1.085908 |
| Critical Value of t | 1.97928 |
| Least Significant Difference | 1.1908 |

Means with the same letter are not significantly different.

| t Grouping |  |  |  | Mean | N | Genotype |
| --- | --- | --- | --- | --- | --- | --- |
|  |  | A |  | 15.0366 | 6 | 18 |
|  |  | A |  |  |  |  |
| B |  | A |  | 14.4751 | 6 | 1 |
| B |  |  |  |  |  |  |
| B |  | C |  | 13.4937 | 6 | 6 |
|  |  | C |  |  |  |  |
| D |  | C |  | 12.9565 | 6 | 3 |
| D |  | C |  |  |  |  |
| D |  | C | E | 12.8580 | 6 | 11 |
| D |  | C | E |  |  |  |
| D | F | C | E | 12.4917 | 6 | 22 |
| D | F | C | E |  |  |  |
| D | F | C | E | 12.4741 | 6 | 4 |
| D | F | C | E |  |  |  |
| D | F | C | E | 12.4712 | 6 | 29 |
| D | F | C | E |  |  |  |
| G | D | F | C | 12.3505 | 6 | 12 |
| G | D | F | C |  |  |  |
| G | D | F | C | 12.3494 | 6 | 10 |
| G | D | F | C |  |  |  |

|  |  |  |  |  |  |  |  |
| --- | --- | --- | --- | --- | --- | --- | --- |
| G | D | F | C | E | 12.3162 | 6 | 17 |
| G | D | F |  | E |  |  |  |
| G | D | F | H | E | 12.1170 | 6 | 16 |
| G | D | F | H | E |  |  |  |
| G | D | F | H | E | 12.0823 | 6 | 19 |
| G | D | F | H | E |  |  |  |
| G | D | F | H | E | 12.0553 | 6 | 5 |
| G | D | F | H | E |  |  |  |
| G | D | F | H | E | 12.0205 | 6 | 9 |
| G | D | F | H | E |  |  |  |
| G | D | F | H | E | 11.9094 | 6 | 23 |
| G | D | F | H | E |  |  |  |
| G | D | F | H | E | 11.8872 | 6 | 27 |
| G | D | F | H | E |  |  |  |
| G | D | F | H | E | 11.8100 | 6 | 31 |
| G |  | F | H | E |  |  |  |
| G | I | F | H | E | 11.7099 | 6 | 8 |
| G | I | F | H | E |  |  |  |

14:00 Thursday, February 18, 2026 45

### The GLM Procedure

#### t Tests (LSD) for Car\_\_mg\_g\_1\_

Means with the same letter are not significantly different.

| t Grouping |  |  |  |  | Mean | N | Genotype |
| --- | --- | --- | --- | --- | --- | --- | --- |
| G | I | F | H | E | 11.6708 | 6 | 13 |
| G | I | F | H |  |  |  |  |
| G | I | F | H |  | 11.6562 | 6 | 20 |
| G | I | F | H |  |  |  |  |
| G | I | F | H |  | 11.5728 | 6 | 7 |
| G | I | F | H |  |  |  |  |
| G | I | F | H | J | 11.4205 | 6 | 15 |
| G | I | F | H | J |  |  |  |
| G | I | F | H | J | 11.4103 | 6 | 21 |
| G | I |  | H | J |  |  |  |
| G | I | K | H | J | 11.1932 | 6 | 25 |
|  | I | K | H | J |  |  |  |
| L | I | K | H | J | 11.0653 | 6 | 14 |
| L | I | K |  | J |  |  |  |
| L | I | K |  | J | 10.5594 | 6 | 30 |
| L | I | K |  | J |  |  |  |
| L | I | K |  | J | 10.5536 | 6 | 28 |
| L |  | K |  | J |  |  |  |
| L |  | K |  | J | 10.3117 | 6 | 26 |
| L |  | K |  |  |  |  |  |
| L |  | K |  |  | 10.1235 | 6 | 2 |
| L |  |  |  |  |  |  |  |
| L |  |  |  |  | 9.9330 | 6 | 24 |

14:00 Thursday, February 18, 2026 46

### The GLM Procedure

#### t Tests (LSD) for H2O2\_\_mmol\_kg\_1\_

NOTE: This test controls the Type I comparisonwise error rate, not the experimentwise error rate.

|  |  |
| --- | --- |
| Alpha | 0.05 |
| Error Degrees of Freedom | 124 |
| Error Mean Square | 0.23834 |
| Critical Value of t | 1.97928 |
| Least Significant Difference | 0.5579 |

Means with the same letter are not significantly different.

| t Grouping | Mean | N | Genotype |
| --- | --- | --- | --- |
| A | 28.0460 | 6 | 5 |
| B | 23.0000 | 6 | 10 |
| C | 21.5824 | 6 | 6 |
| D | 19.2797 | 6 | 3 |
| E | 17.6207 | 6 | 30 |
| F | 16.6398 | 6 | 23 |
| F |  |  |  |
| F | 16.4559 | 6 | 2 |
| F |  |  |  |
| F | 16.2682 | 6 | 4 |
| G |  |  |  |
| G | 13.1686 | 6 | 24 |
| G |  |  |  |
| G | 12.8333 | 6 | 7 |
| H |  |  |  |
| H | 11.5364 | 6 | 13 |
| I |  |  |  |
| I | 10.8889 | 6 | 11 |
| J |  |  |  |
| J | 10.0000 | 6 | 22 |
| J |  |  |  |
| J | 9.6284 | 6 | 1 |
| K |  |  |  |
| K | 8.3678 | 6 | 28 |
| K |  |  |  |
| K | 8.0613 | 6 | 17 |
| K |  |  |  |
| K | 7.8544 | 6 | 20 |
| L |  |  |  |
| L | 7.2338 | 6 | 14 |
| M |  |  |  |
| M | 5.8812 | 6 | 29 |

14:00 Thursday, February 18, 2026 47

###### The GLM Procedure

t Tests (LSD) for H2O2\_\_mmol\_kg\_1\_

Means with the same letter are not significantly different.

| t | Grouping | Mean | N | Genotype |
| --- | --- | --- | --- | --- |
| N | M | 5.4042 | 6 | 27 |
| N |  |  |  |  |
| N |  | 5.3180 | 6 | 21 |
| N |  |  |  |  |
| N | O | 5.0383 | 6 | 12 |
|  | O |  |  |  |
| P | O | 4.6406 | 6 | 25 |
| P |  |  |  |  |
| P | Q | 4.1303 | 6 | 26 |
|  | Q |  |  |  |
| R | Q | 4.0785 | 6 | 8 |
| R | Q |  |  |  |
| R | Q | 4.0460 | 6 | 18 |
| R | Q |  |  |  |
| R | Q | 3.9732 | 6 | 19 |
| R | Q |  |  |  |
| R | Q | 3.8448 | 6 | 9 |
| R |  |  |  |  |
| R |  | 3.5556 | 6 | 31 |
|  | S | 2.3176 | 6 | 16 |
|  | S |  |  |  |
|  | S | 2.2069 | 6 | 15 |

14:00 Thursday, February 18, 2026 48

###### The GLM Procedure

t Tests (LSD) for MDA\_\_mmol\_kg\_1\_

NOTE: This test controls the Type I comparisonwise error rate, not the experimentwise error rate.

|  |  |
| --- | --- |
| Alpha | 0.05 |
| Error Degrees of Freedom | 124 |
| Error Mean Square | 0.033999 |
| Critical Value of t | 1.97928 |
| Least Significant Difference | 0.2107 |

Means with the same letter are not significantly different.

| t Grouping |  |  | Mean | N | Genotype |
| --- | --- | --- | --- | --- | --- |
|  |  | A | 6.5366 | 6 | 24 |
|  |  | B | 4.8382 | 6 | 2 |
|  |  | C | 4.5048 | 6 | 25 |
|  |  | C | 4.4871 | 6 | 1 |
|  |  | C | 4.4468 | 6 | 12 |
| D |  | C | 4.3774 | 6 | 13 |
| D |  | C | 4.3710 | 6 | 22 |
| D |  | C | 4.3183 | 6 | 21 |
| D |  | E | 4.2371 | 6 | 17 |
| D |  | F | 4.1398 | 6 | 10 |
| G |  | F | 4.1274 | 6 | 8 |
| G |  | F | 4.0677 | 6 | 14 |
| G |  | F | 4.0430 | 6 | 7 |
| G | I | F | 4.0263 | 6 | 19 |
| G | I | H | 4.0059 | 6 | 4 |
| G | I | H | 3.9532 | 6 | 6 |
| G | I | H | 3.9097 | 6 | 18 |
| G | I | H | 3.9059 | 6 | 5 |
| K | I | J | 3.8478 | 6 | 30 |
| K | I | J |  |  |  |

14:00 Thursday, February 18, 2026 49

###### The GLM Procedure

t Tests (LSD) for MDA\_\_mmol\_kg\_1\_

Means with the same letter are not significantly different.

| t Grouping |  |  | Mean | N | Genotype |
| --- | --- | --- | --- | --- | --- |
| K | I | J | 3.8468 | 6 | 31 |
| K |  | J | 3.7258 | 6 | 16 |
| K |  | J | 3.7242 | 6 | 27 |
| K |  | L | 3.6640 | 6 | 15 |
| K |  | M | 3.6532 | 6 | 9 |
| K |  | M | 3.5694 | 6 | 3 |
| K |  | M | 3.5522 | 6 | 29 |
| N |  | M | 3.4871 | 6 | 26 |
| N |  | O | 3.2849 | 6 | 20 |
| N |  | O | 3.1398 | 6 | 11 |
| P |  | Q | 3.0376 | 6 | 28 |
| P |  | Q | 2.9226 | 6 | 23 |

14:00 Thursday, February 18, 2026 50

###### The GLM Procedure

t Tests (LSD) for SOD\_\_EU\_g\_1\_Leaf\_

NOTE: This test controls the Type I comparisonwise error rate, not the experimentwise error rate.

Means with the same letter are not significantly different.

| t Grouping |  |  |  |  | Mean | N | Genotype |  |
| --- | --- | --- | --- | --- | --- | --- | --- | --- |
| E |  |  | A |  | 324.061 | 6 | 22 |  |
|  |  |  | A |  |  |  |  |  |
|  | B |  | A |  | 323.689 | 6 | 16 |  |
|  | B |  | A |  |  |  |  |  |
|  | B |  | A | C | 322.000 | 6 | 25 |  |
|  | B |  | A | C |  |  |  |  |
|  | B | D | A | C | 321.456 | 6 | 19 |  |
|  | B | D | A | C |  |  |  |  |
|  | B | D | A | C | 321.144 | 6 | 26 |  |
|  | B | D | A | C |  |  |  |  |
|  | B | D | A | C | 320.950 | 6 | 31 |  |
|  | B | D | A | C |  |  |  |  |
|  | B | D | A | C | F | 319.894 | 6 | 10 |
|  | B | D |  | C | F |  |  |  |
|  | B | D |  | C | F | 319.589 | 6 | 11 |
|  | B | D |  | C | F |  |  |  |
|  | B | D |  | C | F | 319.378 | 6 | 12 |
|  |  | D |  | C | F |  |  |  |
|  |  | D | G | C | F | 317.956 | 6 | 28 |
|  |  | D | G |  | F |  |  |  |
| H | D | G |  | F | 317.528 | 6 | 9 |  |
| H |  | G |  | F |  |  |  |  |
| H |  | G |  | F | 317.039 | 6 | 18 |  |
| H |  | G |  | F |  |  |  |  |
| H |  | G |  | F | 316.889 | 6 | 29 |  |
| H |  | G |  | F |  |  |  |  |
| H |  | G |  | F | 316.478 | 6 | 20 |  |
| H |  | G |  |  |  |  |  |  |
| H |  | G | I |  | 314.578 | 6 | 24 |  |
| H |  | G | I |  |  |  |  |  |
| H | J | G | I |  | 313.944 | 6 | 3 |  |
| H | J | G | I |  |  |  |  |  |
| H | J | G | I |  | 313.933 | 6 | 14 |  |
| H | J |  | I |  |  |  |  |  |
| H | J | K | I |  | 313.489 | 6 | 8 |  |
| H | J | K | I |  |  |  |  |  |
| H | J | K | I |  | 313.428 | 6 | 13 |  |
| H | J | K | I |  |  |  |  |  |

14:00 Thursday, February 18, 2026 51

#### The GLM Procedure

#### t Tests (LSD) for SOD\_EU\_g 1 Leaf\_

Means with the same letter are not significantly different.

| t Grouping |  |  |  | Mean | N | Genotype |
| --- | --- | --- | --- | --- | --- | --- |
| H | J | K | I | 313.178 | 6 | 4 |
|  | J | K | I |  |  |  |
| L | J | K | I | 311.822 | 6 | 2 |
| L | J | K | I |  |  |  |
| L | J | K | I | 310.817 | 6 | 5 |
| L | J | K |  |  |  |  |
| L | J | K | M | 309.711 | 6 | 21 |
| L |  | K | M |  |  |  |
| L |  | K | M | 309.428 | 6 | 1 |
| L |  | K | M |  |  |  |
| L |  | K | M | 309.339 | 6 | 30 |
| L |  |  | M |  |  |  |
| L |  |  | M | 308.128 | 6 | 17 |
| L |  |  | M |  |  |  |
| L |  |  | M | 307.567 | 6 | 15 |
|  |  |  | M |  |  |  |
|  |  |  | M | 306.056 | 6 | 23 |
|  |  | N |  | 298.394 | 6 | 27 |

The GLM Procedure

t Tests (LSD) for CAT\_\_EU\_g\_1\_Leaf\_

NOTE: This test controls the Type I comparisonwise error rate, not the experimentwise error rate.

|  |  |
| --- | --- |
| Alpha | 0.05 |
| Error Degrees of Freedom | 124 |
| Error Mean Square | 41.45282 |
| Critical Value of t | 1.97928 |
| Least Significant Difference | 7.3574 |

Means with the same letter are not significantly different.

| t Grouping |  | Mean | N | Genotype |
| --- | --- | --- | --- | --- |
|  | A | 135.366 | 6 | 12 |
|  | B | 125.428 | 6 | 28 |
|  | B | 124.638 | 6 | 27 |
| C | B | 123.761 | 6 | 25 |
| C | B | 122.669 | 6 | 7 |
| C | D | 117.947 | 6 | 4 |
| C | D | 114.398 | 6 | 8 |
| E | D | 112.475 | 6 | 20 |
| E | F | 110.130 | 6 | 24 |
| E | F | 109.539 | 6 | 16 |
| E | F | 109.456 | 6 | 23 |
| E | F | 108.487 | 6 | 6 |
|  | F | 104.819 | 6 | 5 |
| H | F | 103.157 | 6 | 13 |
| H | I | 101.867 | 6 | 2 |
| H | I | 101.760 | 6 | 29 |
| H | I | 101.160 | 6 | 3 |
| H | I | 100.857 | 6 | 26 |
| H | I | 100.820 | 6 | 11 |
| H | I |  |  |  |

The GLM Procedure

t Tests (LSD) for CAT\_\_EU\_g\_1\_Leaf\_

Means with the same letter are not significantly different.

| t Grouping |  | Mean | N | Genotype |
| --- | --- | --- | --- | --- |
| H | I | 100.672 | 6 | 1 |
| H | I |  |  |  |
| H | I | 98.742 | 6 | 17 |
|  | I |  |  |  |
|  | I | 96.651 | 6 | 9 |
|  | J |  |  |  |
| K | J | 92.925 | 6 | 22 |
| K | J |  |  |  |
| K | J | 92.423 | 6 | 14 |
| K | J | 92.310 | 6 | 15 |
| K |  |  |  |  |
| K |  |  |  |  |
| K | L | 88.978 | 6 | 21 |
|  | L |  |  |  |
|  | L | 83.878 | 6 | 30 |
|  | M | 75.349 | 6 | 10 |
|  | M |  |  |  |
| N | M | 69.058 | 6 | 31 |
| N |  |  |  |  |
| N |  | 67.376 | 6 | 18 |
| N |  |  |  |  |
| N |  | 63.064 | 6 | 19 |

14:00 Thursday, February 18, 2026 54

The GLM Procedure

t Tests (LSD) for POD\_\_EU\_g\_1\_Leaf\_

NOTE: This test controls the Type I comparisonwise error rate, not the experimentwise error rate.

|  |  |
| --- | --- |
| Alpha | 0.05 |
| Error Degrees of Freedom | 124 |
| Error Mean Square | 33308.78 |
| Critical Value of t | 1.97928 |
| Least Significant Difference | 208.56 |

Means with the same letter are not significantly different.

| t | Grouping | Mean | N | Genotype |
| --- | --- | --- | --- | --- |
|  | A | 19552.3 | 6 | 19 |
|  | B | 17763.4 | 6 | 18 |
|  | C | 11107.8 | 6 | 6 |
|  | D | 9705.9 | 6 | 17 |
|  | E | 7384.2 | 6 | 5 |
|  | F | 6726.5 | 6 | 3 |
|  | G | 6495.1 | 6 | 28 |
|  | H | 5190.3 | 6 | 13 |
|  | H |  |  |  |
|  | H | 5048.7 | 6 | 1 |
|  | I | 4781.0 | 6 | 2 |
|  | J | 3828.8 | 6 | 22 |
|  | J |  |  |  |
| K | J | 3756.1 | 6 | 26 |
| K |  |  |  |  |
| K |  | 3590.2 | 6 | 21 |
| K |  |  |  |  |
| K |  | 3581.7 | 6 | 24 |
| K |  |  |  |  |
| K |  | 3558.8 | 6 | 15 |
|  | L | 3130.7 | 6 | 31 |
|  | M | 2774.0 | 6 | 4 |
|  | M |  |  |  |
|  | M | 2570.9 | 6 | 30 |
|  | N | 2277.8 | 6 | 11 |
|  | N |  |  |  |

14:00 Thursday, February 18, 2026 55

The GLM Procedure

t Tests (LSD) for POD\_\_EU\_g\_1\_Leaf\_

Means with the same letter are not significantly different.

| t | Grouping | Mean | N | Genotype |
| --- | --- | --- | --- | --- |
| 0 | N | 2074.5 | 6 | 27 |
| 0 |  |  |  |  |
| 0 | P | 1884.3 | 6 | 16 |
|  | P |  |  |  |
| Q | P | 1796.1 | 6 | 10 |
| Q | P |  |  |  |
| Q | P | 1725.0 | 6 | 23 |
| Q |  |  |  |  |
| Q |  | 1638.8 | 6 | 12 |
| Q |  |  |  |  |
| Q |  | 1612.4 | 6 | 20 |
|  | R | 1350.7 | 6 | 25 |
|  | R |  |  |  |
| S | R | 1238.9 | 6 | 29 |
| S |  |  |  |  |
| S | T | 1137.3 | 6 | 14 |
|  | T |  |  |  |
|  | T | 978.0 | 6 | 9 |
|  | T |  |  |  |
|  | T | 936.2 | 6 | 7 |
|  | T |  |  |  |
|  | T | 931.0 | 6 | 8 |

14:00 Thursday, February 18, 2026 56

The GLM Procedure

t Tests (LSD) for PH\_cm\_

NOTE: This test controls the Type I comparisonwise error rate, not the experimentwise error rate.

|  |  |
| --- | --- |
| Alpha | 0.05 |
| Error Degrees of Freedom | 124 |
| Error Mean Square | 0.928538 |
| Critical Value of t | 1.97928 |
| Least Significant Difference | 0.2797 |

Means with the same letter are not significantly different.

| t | Grouping | Mean | N | FC____ |
| --- | --- | --- | --- | --- |
|  | A | 27.8315 | 93 | 100 |
|  | B | 21.7803 | 93 | 60 |

14:00 Thursday, February 18, 2026 57

The GLM Procedure

t Tests (LSD) for SD\_cm\_

NOTE: This test controls the Type I comparisonwise error rate, not the experimentwise error rate.

|  |  |
| --- | --- |
| Alpha | 0.05 |
| Error Degrees of Freedom | 124 |
| Error Mean Square | 0.004087 |
| Critical Value of t | 1.97928 |
| Least Significant Difference | 0.0186 |

Means with the same letter are not significantly different.

| t | Grouping | Mean | N | FC____ |
| --- | --- | --- | --- | --- |
|  | A | 2.255054 | 93 | 100 |
|  | B | 1.846344 | 93 | 60 |

14:00 Thursday, February 18, 2026 58

The GLM Procedure

t Tests (LSD) for LA\_cm2\_

NOTE: This test controls the Type I comparisonwise error rate, not the experimentwise error rate.

|  |  |
| --- | --- |
| Alpha | 0.05 |
| Error Degrees of Freedom | 124 |
| Error Mean Square | 268.3265 |
| Critical Value of t | 1.97928 |
| Least Significant Difference | 4.7546 |

Means with the same letter are not significantly different.

| t Grouping | Mean | N | FC____ |
| --- | --- | --- | --- |
| A | 414.551 | 93 | 100 |
| B | 253.041 | 93 | 60 |

14:00 Thursday, February 18, 2026 59

The GLM Procedure

t Tests (LSD) for PFW\_\_g\_\_

NOTE: This test controls the Type I comparisonwise error rate, not the experimentwise error rate.

|  |  |
| --- | --- |
| Alpha | 0.05 |
| Error Degrees of Freedom | 124 |
| Error Mean Square | 0.460161 |
| Critical Value of t | 1.97928 |
| Least Significant Difference | 0.1969 |

Means with the same letter are not significantly different.

| t Grouping | Mean | N | FC____ |
| --- | --- | --- | --- |
| A | 20.09892 | 93 | 100 |
| B | 12.56129 | 93 | 60 |

14:00 Thursday, February 18, 2026 60

The GLM Procedure

t Tests (LSD) for RFW\_\_g\_\_

NOTE: This test controls the Type I comparisonwise error rate, not the experimentwise error rate.

|  |  |
| --- | --- |
| Alpha | 0.05 |
| Error Degrees of Freedom | 124 |
| Error Mean Square | 0.398503 |
| Critical Value of t | 1.97928 |
| Least Significant Difference | 0.1832 |

Means with the same letter are not significantly different.

| t Grouping | Mean | N | FC____ |
| --- | --- | --- | --- |
| A | 7.22527 | 93 | 100 |
| B | 4.30376 | 93 | 60 |

14:00 Thursday, February 18, 2026 61

The GLM Procedure

t Tests (LSD) for PDW\_\_g\_\_

NOTE: This test controls the Type I comparisonwise error rate, not the experimentwise error rate.

|  |  |
| --- | --- |
| Alpha | 0.05 |
| Error Degrees of Freedom | 124 |
| Error Mean Square | 0.009234 |
| Critical Value of t | 1.97928 |
| Least Significant Difference | 0.0279 |

Means with the same letter are not significantly different.

t Grouping Mean N FC\_\_\_\_\_

A 1.79215 93 100

B 0.97129 93 60

14:00 Thursday, February 18, 2026 62

The GLM Procedure

t Tests (LSD) for RDW\_\_g\_

NOTE: This test controls the Type I comparisonwise error rate, not the experimentwise error rate.

Alpha 0.05
Error Degrees of Freedom 124
Error Mean Square 0.00157
Critical Value of t 1.97928
Least Significant Difference 0.0115

Means with the same letter are not significantly different.

t Grouping Mean N FC\_\_\_\_\_

A 0.459785 93 100

B 0.247849 93 60

14:00 Thursday, February 18, 2026 63

The GLM Procedure

t Tests (LSD) for EC\_\_\_\_\_

NOTE: This test controls the Type I comparisonwise error rate, not the experimentwise error rate.

Alpha 0.05
Error Degrees of Freedom 124
Error Mean Square 12.79635
Critical Value of t 1.97928
Least Significant Difference 1.0383

Means with the same letter are not significantly different.

t Grouping Mean N FC\_\_\_\_\_

A 41.5762 93 60

B 21.9904 93 100

14:00 Thursday, February 18, 2026 64

The GLM Procedure

t Tests (LSD) for LRWC\_\_\_\_\_

NOTE: This test controls the Type I comparisonwise error rate, not the experimentwise error rate.

Alpha 0.05
Error Degrees of Freedom 124
Error Mean Square 33.11233
Critical Value of t 1.97928
Least Significant Difference 1.6702

Means with the same letter are not significantly different.

t Grouping Mean N FC\_\_\_\_\_

A 71.0625 93 100

B 62.2783 93 60

14:00 Thursday, February 18, 2026 65

The GLM Procedure

t Tests (LSD) for Chla\_\_mg\_g\_1\_

NOTE: This test controls the Type I comparisonwise error rate, not the experimentwise error rate.

|  |  |
| --- | --- |
| Alpha | 0.05 |
| Error Degrees of Freedom | 124 |
| Error Mean Square | 0.120174 |
| Critical Value of t | 1.97928 |
| Least Significant Difference | 0.1006 |

Means with the same letter are not significantly different.

| t Grouping | Mean | N | FC____ |
| --- | --- | --- | --- |
| A | 5.65378 | 93 | 100 |
| B | 3.31452 | 93 | 60 |

14:00 Thursday, February 18, 2026 66

The GLM Procedure

t Tests (LSD) for Chlb\_\_mg\_g\_1\_

NOTE: This test controls the Type I comparisonwise error rate, not the experimentwise error rate.

|  |  |
| --- | --- |
| Alpha | 0.05 |
| Error Degrees of Freedom | 124 |
| Error Mean Square | 0.025918 |
| Critical Value of t | 1.97928 |
| Least Significant Difference | 0.0467 |

Means with the same letter are not significantly different.

| t Grouping | Mean | N | FC____ |
| --- | --- | --- | --- |
| A | 3.50249 | 93 | 100 |
| B | 1.74891 | 93 | 60 |

14:00 Thursday, February 18, 2026 67

The GLM Procedure

t Tests (LSD) for Total\_Ch1\_\_mg\_g\_1\_

NOTE: This test controls the Type I comparisonwise error rate, not the experimentwise error rate.

|  |  |
| --- | --- |
| Alpha | 0.05 |
| Error Degrees of Freedom | 124 |
| Error Mean Square | 0.170501 |
| Critical Value of t | 1.97928 |
| Least Significant Difference | 0.1199 |

Means with the same letter are not significantly different.

| t Grouping | Mean | N | FC____ |
| --- | --- | --- | --- |
| A | 9.15454 | 93 | 100 |
| B | 5.06288 | 93 | 60 |

14:00 Thursday, February 18, 2026 68

The GLM Procedure

t Tests (LSD) for Car\_\_mg\_g\_1\_

NOTE: This test controls the Type I comparisonwise error rate, not the experimentwise error rate.

|  |  |
| --- | --- |
| Alpha | 0.05 |
| Error Degrees of Freedom | 124 |
| Error Mean Square | 1.085908 |
| Critical Value of t | 1.97928 |
| Least Significant Difference | 0.3025 |

Means with the same letter are not significantly different.

| t Grouping | Mean | N | FC____ |
| --- | --- | --- | --- |
| A | 14.1222 | 93 | 100 |
| B | 9.7704 | 93 | 60 |

14:00 Thursday, February 18, 2026 69

The GLM Procedure

t Tests (LSD) for H2O2\_\_mmol\_kg\_1\_

NOTE: This test controls the Type I comparisonwise error rate, not the experimentwise error rate.

|  |  |
| --- | --- |
| Alpha | 0.05 |
| Error Degrees of Freedom | 124 |
| Error Mean Square | 0.23834 |
| Critical Value of t | 1.97928 |
| Least Significant Difference | 0.1417 |

Means with the same letter are not significantly different.

| t Grouping | Mean | N | FC____ |
| --- | --- | --- | --- |
| A | 12.82974 | 93 | 60 |
| B | 7.35741 | 93 | 100 |

14:00 Thursday, February 18, 2026 70

The GLM Procedure

t Tests (LSD) for MDA\_\_mmol\_kg\_1\_

NOTE: This test controls the Type I comparisonwise error rate, not the experimentwise error rate.

|  |  |
| --- | --- |
| Alpha | 0.05 |
| Error Degrees of Freedom | 124 |
| Error Mean Square | 0.033999 |
| Critical Value of t | 1.97928 |
| Least Significant Difference | 0.0535 |

Means with the same letter are not significantly different.

| t Grouping | Mean | N | FC____ |
| --- | --- | --- | --- |
| A | 5.05730 | 93 | 60 |
| B | 2.92667 | 93 | 100 |

14:00 Thursday, February 18, 2026 71

The GLM Procedure

t Tests (LSD) for SOD\_\_EU\_g\_1\_Leaf\_

NOTE: This test controls the Type I comparisonwise error rate, not the experimentwise error rate.

|  |  |
| --- | --- |
| Alpha | 0.05 |
| Error Degrees of Freedom | 124 |
| Error Mean Square | 14.62488 |
| Critical Value of t | 1.97928 |
| Least Significant Difference | 1.11 |

Means with the same letter are not significantly different.

| t Grouping | Mean | N | FC____ |
| --- | --- | --- | --- |
| A | 330.3989 | 93 | 60 |
| B | 296.2595 | 93 | 100 |

14:00 Thursday, February 18, 2026 72

The GLM Procedure

t Tests (LSD) for CAT\_\_EU\_g\_1\_Leaf\_

NOTE: This test controls the Type I comparisonwise error rate, not the experimentwise error rate.

|  |  |
| --- | --- |
| Alpha | 0.05 |
| Error Degrees of Freedom | 124 |
| Error Mean Square | 41.45282 |
| Critical Value of t | 1.97928 |
| Least Significant Difference | 1.8688 |

Means with the same letter are not significantly different.

| t Grouping | Mean | N | FC____ |
| --- | --- | --- | --- |
| A | 124.6595 | 93 | 60 |
| B | 78.5768 | 93 | 100 |

14:00 Thursday, February 18, 2026 73

The GLM Procedure

t Tests (LSD) for POD\_\_EU\_g\_1\_Leaf\_

NOTE: This test controls the Type I comparisonwise error rate, not the experimentwise error rate.

|  |  |
| --- | --- |
| Alpha | 0.05 |
| Error Degrees of Freedom | 124 |
| Error Mean Square | 33308.78 |
| Critical Value of t | 1.97928 |
| Least Significant Difference | 52.974 |

Means with the same letter are not significantly different.

| t Grouping | Mean | N | FC____ |
| --- | --- | --- | --- |
| A | 7004.70 | 93 | 60 |
| B | 2035.78 | 93 | 100 |
